# Pleistocene climatic oscillations impact the diversification of deer mice (*Peromyscus maniculatus)* and the independent evolution of ecotypes

**DOI:** 10.64898/2026.01.26.699144

**Authors:** Robert A. Boria, T. Brock Wooldridge, Andreas F. Kautt, Kemi Ashing-Giwa, Sade McFadden, Christopher Kirby, Scott Edwards, Hopi E. Hoekstra

**Affiliations:** Department of Organismic and Evolutionary Biology, Harvard University, Cambridge, MA 02138; Museum of Comparative Zoology, Harvard University, Cambridge, MA 02138; Department of Molecular and Cellular Biology Harvard University, Cambridge, MA 02138; Department of Biology, San Francisco State University, San Francisco, CA 942138; Ecology and Evolutionary Biology Department, University of California Santa Cruz, Santa Cruz, CA 95064; Department of Biology, Washington University in St. Louis, St. Louis, MO 63130

## Abstract

A central question in evolutionary biology is whether local adaptation is predictable when a species repeatedly encounters similar environments. The deer mouse, *Peromyscus maniculatus*, has a range of over 13 million km^2^ in North America and may be found in nearly every terrestrial habitat. Because of their abundance and wide habitat preference, deer mice and closely related *Peromyscus*, which we refer to as the *P. maniculatus* species complex, are at the forefront of studies of biogeography and local adaptation. Here, we undertake a comprehensive survey of genome-wide and phenotypic diversity to characterize the recent evolutionary history of this group. We sequenced whole genomes from 232 individuals across their range, representing the most thorough genetic sampling of the *P. maniculatus* species complex to date. We identify six geographically delineated clades, several of which encompass both classically recognized *P. maniculatus* subspecies as well as other recognized species. Ecological niche modelling suggests that this geographic structure resulted from rapid post-LGM range expansion and adaptation to emerging habitats. Our morphological measurements of 979 specimens and field data compiled from over 28,000 museum records show that deer mice in forests across the range consistently have longer tails, larger feet, bigger ears, and elongated whiskers. These traits constitute an arboreal ecotype that has evolved at least three times independently, and was likely lost in other parts of the range as populations moved out of forested habitat. Altogether, these results suggest that post-LGM increases in forested habitat drove the parallel evolution of arboreal ecotypes across the deer mouse range.

## Introduction

Forecasting future biodiversity under natural and anthropogenic impacts is a central goal in ecology and evolutionary biology (Dawson et al., 2011). Crucial to this is understanding how intraspecific genetic diversity is spatially distributed (Yannic et al., 2014). Climate change can modify the distribution of genetic diversity through several mechanisms, including range shifts of historically significant lineages and local adaptation to emerging environmental conditions (Pauls et al., 2013). All are frequently associated with reductions in overall genetic diversity (Bálint et al., 2011; Pironon et al., 2016). Many temperate species have already responded to recent climate change, and others have shown a decline in intraspecific genetic diversity extending back to the Last Glacial Maximum (LGM), 21,000 years ago (Hewitt, 2000; Magyari et al., 2011; Miraldo et al., 2016).

The Pleistocene glacial-interglacial cycles have been recognized as one of the primary drivers of intraspecific variation across the tree of life (Hewitt 2000, 2004). These cycles, occurring from 1.5 MYA to 21 KYA, were a succession of cool glacial periods followed by warm interglacial intervals, with short transitions between (Dansgaard *et al*. 1993). For example, during the Last Glacial Maximum (LGM) enormous ice sheets covered most of the northern portion of the continent. Climatic conditions south of the ice sheets were generally drier and cooler than at present (Hopkins 2013), except in the southwest where wetter conditions prevailed (Clark *et al*. 2012). During the LGM, species in North America largely experienced a reduction in habitat due to the spread of continental ice sheets, environmental conditions outside of their physiological range, and/or fragmentation of primary habitat (Hewitt 2004; Nogués-Bravo *et al*. 2008). For species with large geographic distributions, glacial-interglacial cycles could lead to repeated expansions and contractions into and out of ecological niches, in turn repeatedly selecting for the same sets of traits (i.e., ecotypes) from a species’ standing variation (Nevado *et al*. 2018; Sepúlveda-Espinoza *et al*. 2022).

The enormous diversity of the deer mouse, *Peromyscus maniculatus* (Wagner, 1845), may be a direct result of Pleistocene glacial cycles. *P. maniculatus* is indigenous to and distributed widely across North America and can be found in nearly every terrestrial habitat (Shorter *et al*. 2012; W.H Osgood, 1909). Mitochondrial DNA evidence suggests that *P. maniculatus* consists of at least six distinct geographic clades (Dragoo *et al*. 2006; Kalkvik *et al*. 2012; Natarajan *et al*. 2015), and some of these mitochondrial clades have been proposed as distinct species (Greenbaum *et al*. 2017; Bradley *et al*. 2019; Greenbaum, Honeycutt and Chirhart 2019). There are also morphological and regional forms that have received their own species designations: *P*. *arcticus* (Osgood 1909; Lucid and Cook 2007; *P. keeni* (Rhoads 1895; Zheng, Arbogast and Kenagy 2003), *P. labecula* (Elliot 1903), *P. melanotis* (Allen and Chapman 1897); and *P. polionotus (Osgood 1909; Hall 1981)*, Bradley et al. 2019; Greenbaum, Honeycutt and Chirhart 2019). The phylogenetic placement of these lineages is still under scrutiny (Dragoo *et al*. 2006; Kalkvik *et al*. 2012; Natarajan *et al*. 2015; Sawyer *et al*. 2017; Bradley *et al*. 2019; Boria and Blois 2023). Given this taxonomic complexity, we will refer to all sampled groups as populations of *P. maniculatus* or subspecies (i.e. *P. m. maniculatus*, *P. m. sonoriensis*) for the purposes of this work. We refer to this set of populations and subspecies as the *P. maniculatus* “species complex”.

Despite this diversity of labels, one can see striking morphological similarity between deer mice populations at opposite ends of the continent. Over a century ago, natural historians began noting consistent differences between deer mice inhabiting the forests and prairies of North America (Osgood 1909). In particular, forest mice tended to have longer tails than prairie mice, even when the habitats were directly adjacent and migration could be expected. Decades of work following this showed that deer mouse populations across the species’ range could often be divided into two groups: one with long tails, feet, and ears occupying forests, and one with shorter tails, feet, and ears occupying prairies or similarly open habitats (King 1968). These trait differences are thought to be driven by selection for arboreality in forests, as evidenced by other arboreal rodents (Layne 1970; Samuels and Van Valkenburgh 2008; Verde Arregoitia, Fisher and Schweizer, Manuel 2017; Mincer and Russo 2020). Therefore, we refer to the correlated set of traits seen in forested habitats as the “arboreal” ecotype. While the arboreal ecotype and its counterpart the “prairie” ecotype, have been described in detail for select populations, it remains to be seen how widespread these adaptations are. Other traits, for example whisker length, are associated with arboreal deer mice but are far less studied (Gable 2020; Grant and Goss 2022). It’s thought that the longer tails, larger ears, larger hind feet, and larger whiskers associated with tree-dwellers all aid in navigating a dense, complex environment.

To better understand how the Pleistocene glacial cycles influenced the diversification of deer mice and led to the repeated evolution of arboreal and prairie ecotypes, we perform extensive whole genome re-sequencing and phenotyping from across the *P. maniculatus* range. Despite this sampling breadth, we consistently recover six lineages across population structure and phylogenetic analyses. Next, we identify and delimit suitable conditions across the known range of *P. maniculatus* under current climatic conditions and reveal a substantial reduction of suitable habitat during the LGM. We predict that range expansion and contraction during the Pleistocene glacial cycles, particularly after the LGM, drove repeated phenotypic evolution as deer mice dispersed to newly available habitats. Consistent with this hypothesis we observe the evolution of arboreal ecotypes in at least three lineages across North America, along with several instances of partial ecotype evolution. We find that many of these arboreal populations are genetically and geographically close to prairie-ecotype deer mice, suggesting that divergent selection to maintain trait differences could be widespread. In sum, these findings detail a history of widespread and rapid local adaptation by deer mouse populations across the continent following the last glacial maximum.

## Results

### Whole genome sequencing of *P. maniculatus* and related groups

We generated whole genome sequence data from 232 individuals representing *P. maniculatus* and groups that have previously been referred to as *P. maniculatus*, including *P. arcticus*, *P. gambelli*, *P. keeni*, *P. labecula*, *P. melanotis, P. polionotus*, and *P. sonoriensis* (Bradley et al. 2019; Greenbaum, Honeycutt and Chirhart 2019). The sequenced samples span North America and encompass all ends of the reported *P. maniculatus* range (Fig. 1). Whole-genome sequence coverage ranged from 5X to 34X and estimates of individual-level heterozygosity ranged from 0.31% to 0.47%. Unless otherwise noted, all downstream analyses were based on a reduced autosomal call set in which we removed any variants other than bi-allelic SNPs, removed sites with more than 5% missing data, and randomly sampled one variant every 10 kb to account for non-independence of physically linked sites. The final call set comprised 170,510 SNPs.

**Figure 1.**
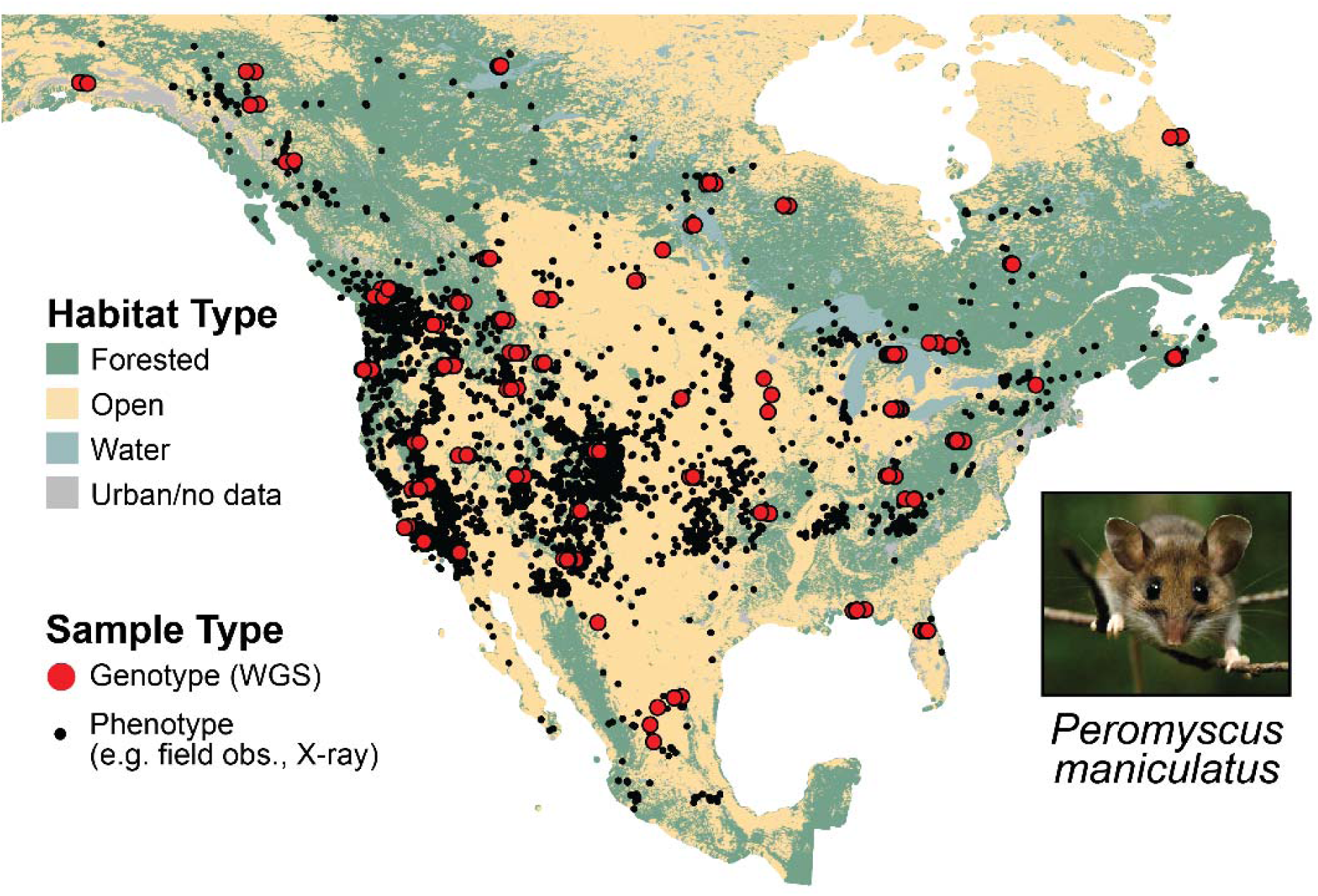
Distribution of samples used in this study. Each point represents an individual from the *Peromyscus maniculatus* species complex. Photo by Phil Myers, licensed under CC BY-NC-SA 3.0.

### Population structure analyses recover geographically distributed groups

Population structure was consistent across both non-spatial and spatially explicit analyses (Fig. S1). First, discriminant analysis of principal components (*DAPC*; non-spatial) (Jombart 2008; Jombart and Ahmed 2011) recovered *K* = 7 populations with the lowest BIC score (Figure S1A; Table S2). Complementary to this, *ADMIXTURE* (non-spatial; Alexander, Novembre and Lange 2009) cross-validation indicated a *K* = 7 model best fit the data (Fig. S1B; Table S3). We recovered the same lineages using a spatially explicit population structure inference, *TESS3r* (incorporates geography and genotypic data; Caye *et al*. 2016, 2018), as we recovered with the non-spatial methods (Fig. S1). *TESS3r* exhibited the lowest cross-validation score at K=10, although *K* = 6-10 were separated by < 0.004 (Fig. S3). The geographic structure recovered by *TESS3r* aligns with mitochondrial and geographic groups previously identified as *P. m. keeni, P. m. maniculatus*, *P. m. melanotis*, *P. m. polionotus*, and *P. m. sonoriensis*, (Fig. 2), as well as one newly identified group: a midwestern population (*P. m.* Midwest hereafter). While we recover finer population structure with increasing K for all methods, clustering is most similar across methods at K = 6. Highly admixed individuals identified in these analyses were removed to generate an additional ‘reduced’ dataset (n=211) for subsequent analyses (Fig. S2; Table S4).

**Figure 2.**
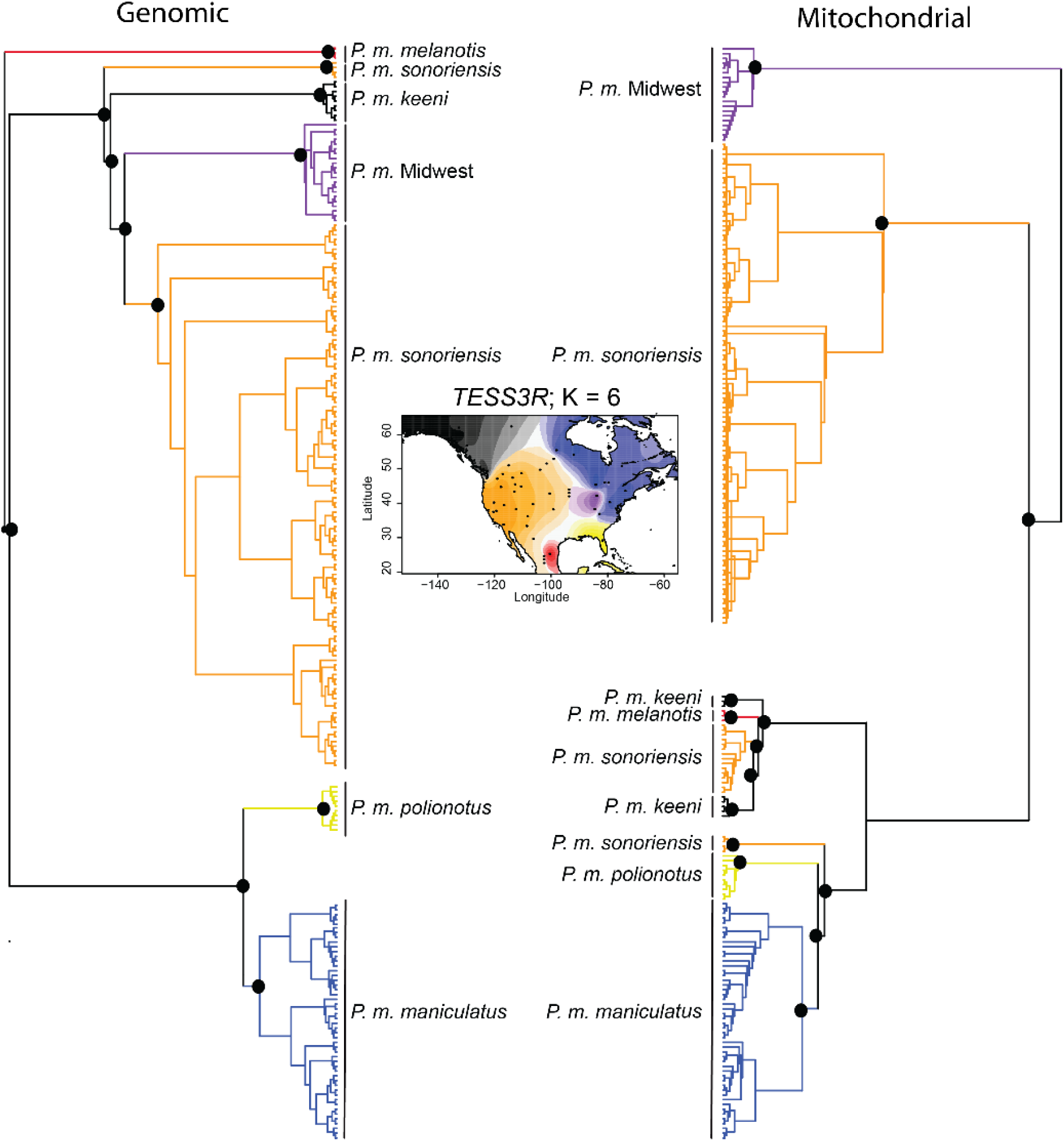
Maximum-likelihood phylogenies based on genomic SNPs (left) and mitochondrial SNPs (right). Trees are rooted with *P. leucopus* (not shown). Clade coloring on trees corresponds to spatial coloring in map inset, which represents the TESS3R spatially explicit population structure at K = 6. Additional TESS3R results are Fig. S2. Black dots indicate greater than 70% node support, and are only displayed for select deeper splits to reduce clutter.

### Whole-genome phylogenetics agrees with population genomic structure

Multiple whole-genome phylogenetic reconstruction approaches confirm the population structure results. Both the maximum likelihood (IQTree; Minh et al. 2020) and coalescent trees (SVDQuartets; Chifman and Kubatko 2014), recover six distinct clades. All six of these clades match the six groups consistently recovered by all population structure approaches and include *P. m. keeni*, *P. m. maniculatus*, *P. m. polionotus*, *P. m. melanotis*, *P. m.* Midwest, and *P. m. sonoriensis* (Fig. 2). Besides these six clades, the primary feature of both autosomal phylogenies is a clear east-west split roughly corresponding to the Mississippi River (Fig. 2; Fig. S4; Fig. S5). While the mitochondrial tree recovers some of these same groupings, it also exhibits a deep, highly supported split that does not correspond to a geographic boundary. Two groups recovered by the population structure analyses and autosomal phylogenies, *P. m. sonoriensis* and *P. m. keeni*, are split and become paraphyletic in the mitochondrial tree. In contrast to its position outside all other deer mouse diversity in the autosomal trees, *P. m. melanotis* is nested within deer mouse mitochondrial diversity. In general, the clustering of individuals shown by the mitochondrial tree is at odds with relationships that might be expected from geographic proximity or our population structure results.

We detected evidence for significant gene flow between *P. m. maniculatus* and *P. m. sonoriensis*. Introgression analysis using *Dsuite* (Patterson et al. 2012; Malinsky et al. 2020) showed that 8/20 trios had significant D-statistics and f_4_-ratios. (post-Benjamini-Hochberg correction; Table S5). Of these eight, seven included either or both *P. m. maniculatus* or *P. m. sonoriensis*. This result was corroborated by the f-branch statistic, indicating that the gene flow estimated here is likely driven by *P. m. maniculatus*, *P. m. sonoriensis*, and to a lesser extent *P. m.* Midwest (Fig. S6).

### Species distribution models support rapid expansion of deer mouse range following the last glacial maximum

Ecological niche modelling (ENM) of *P. maniculatus*, using Maxent (Phillips, Anderson and Schapir 2006; Phillips *et al*. 2017), indicates a significant increase in suitable habitat following the last glacial maximum, ∼19 thousand years before present (Fig. 3; Table S6). Beginning at the present day, we find that ENMs based on climate models confirm suitable habitat for deer mice across the known range (Fig. 3A). At 10 KYA the northern limit of suitability begins to retreat (Supplementary Video). By 19 KYA, when the Laurentide ice sheet was last at its largest, suitable habitat fell below the 40th parallel (Fig. 3B). In contrast, the southern range stays consistently habitable over these glacial cycles. Our analysis of effective population size shows that the primary deer mouse lineages indeed began to diversify around and after the LGM, in particular *P. m. sonoriensis* (Fig. S7), which is consistent with a negative Tajima’ D value indicating a post-LGM expansion (Fig. S8) This suggests that deer mice rapidly expanded northward from a southern refuge following the retreat of the Laurentide ice sheet.

**Figure 3.**
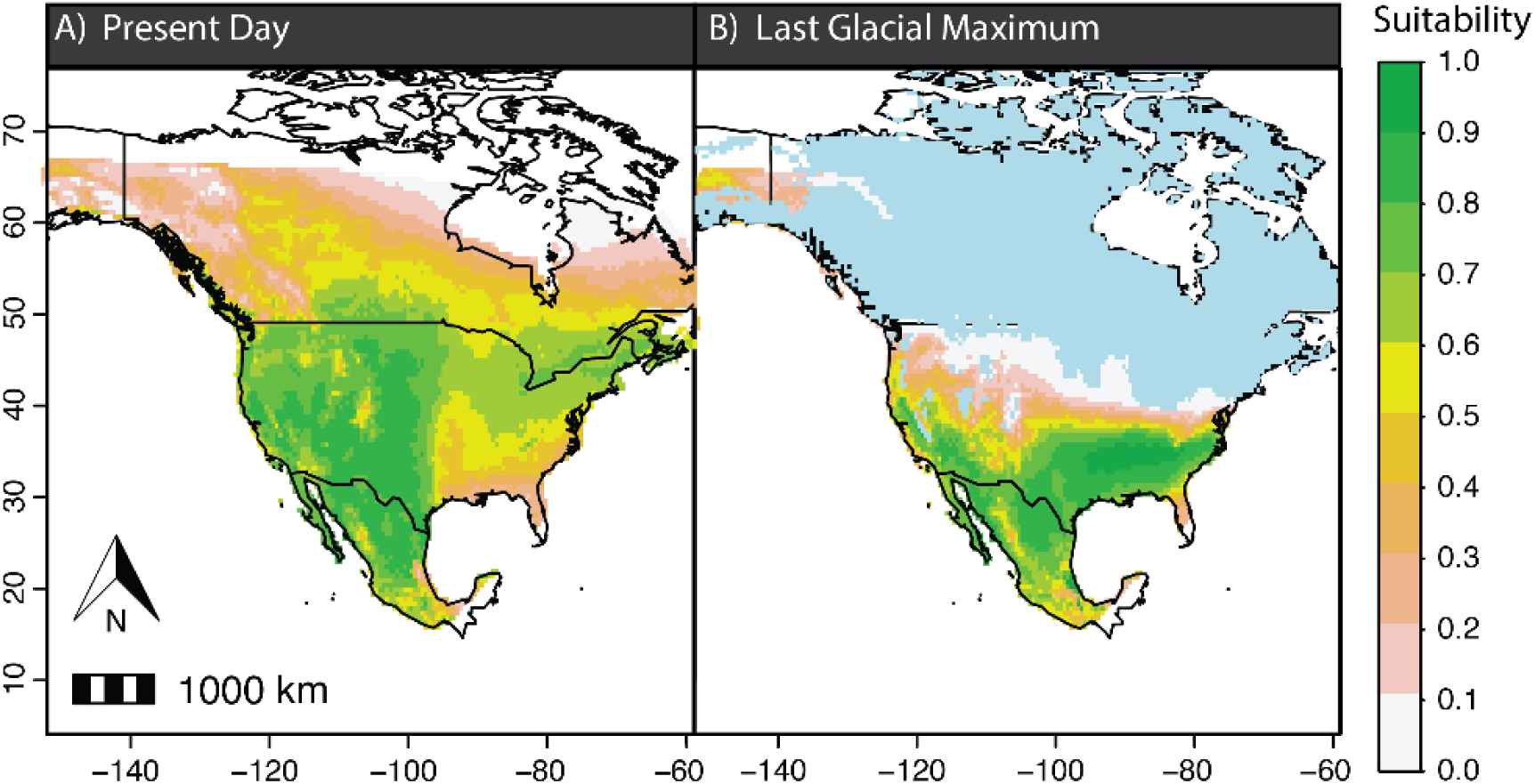
Ecological niche models of *P. maniculatus* throughout its range in North America: A) inferred habitat suitability at the present; B) inferred habitat suitability during the Last Glacial Maximum (LGM). Note the reduction in suitable habitat during the LGM and expansion during the present climatic conditions.

### Range-wide associations between habitat and morphology

Data contained in 28,883 museum records shows that deer mouse morphology is significantly associated with habitat type across the species’ range. To determine this, we first classified habitat as ‘forest’ (e.g. needleleaf forest) or ‘open’ (e.g. grassland) at 1-km resolution following Kingsley et al. (2017). We then analyzed associations between habitat type and three traits common in mammalogy collections - tail, foot, and ear length - for *P. maniculatus* records dating back to 1818. We find significantly longer tails, feet, and ears in mice collected from forest habitats as opposed to open habitats after controlling for body size (Fig. S9, two-sided t-tests: p < 2.2e-16). Of these traits, tail length shows the most dramatic difference - for a deer mouse of mean body size (x= 85 mm), the tail of a forest habitat mouse is on average 15% longer (10 mm) than the tail of an open habitat mouse.

### Repeated independent evolution of arboreal and prairie ecotypes

X-ray imaging and external phenotyping of 979 deer mouse specimens from across North America (Table S7) allowed us to classify populations by ecotype. For this, we focused on skeletal traits (e.g. caudal vertebra number) and external phenotypes including whisker length (Fig. 4A; Table S7). We performed linear discriminant analysis (LDA) using these data from four known arboreal and prairie populations as training data to create a model classifying ecotypes with 90.8% accuracy (Fig. S10). Our classification results show that arboreal populations may be found as far apart as 3,910 km, and prairie populations as far apart as 4,929 km (Fig. 4B; Table S8). For reference, the maximum distance between any two points in the IUCN-estimated *P. maniculatus* range is 5,362 km. We also observe several populations classified as ‘intermediate’ ecotypes inhabiting forested and/or alpine areas across the species range (Fig. 4C).

**Figure 4.**
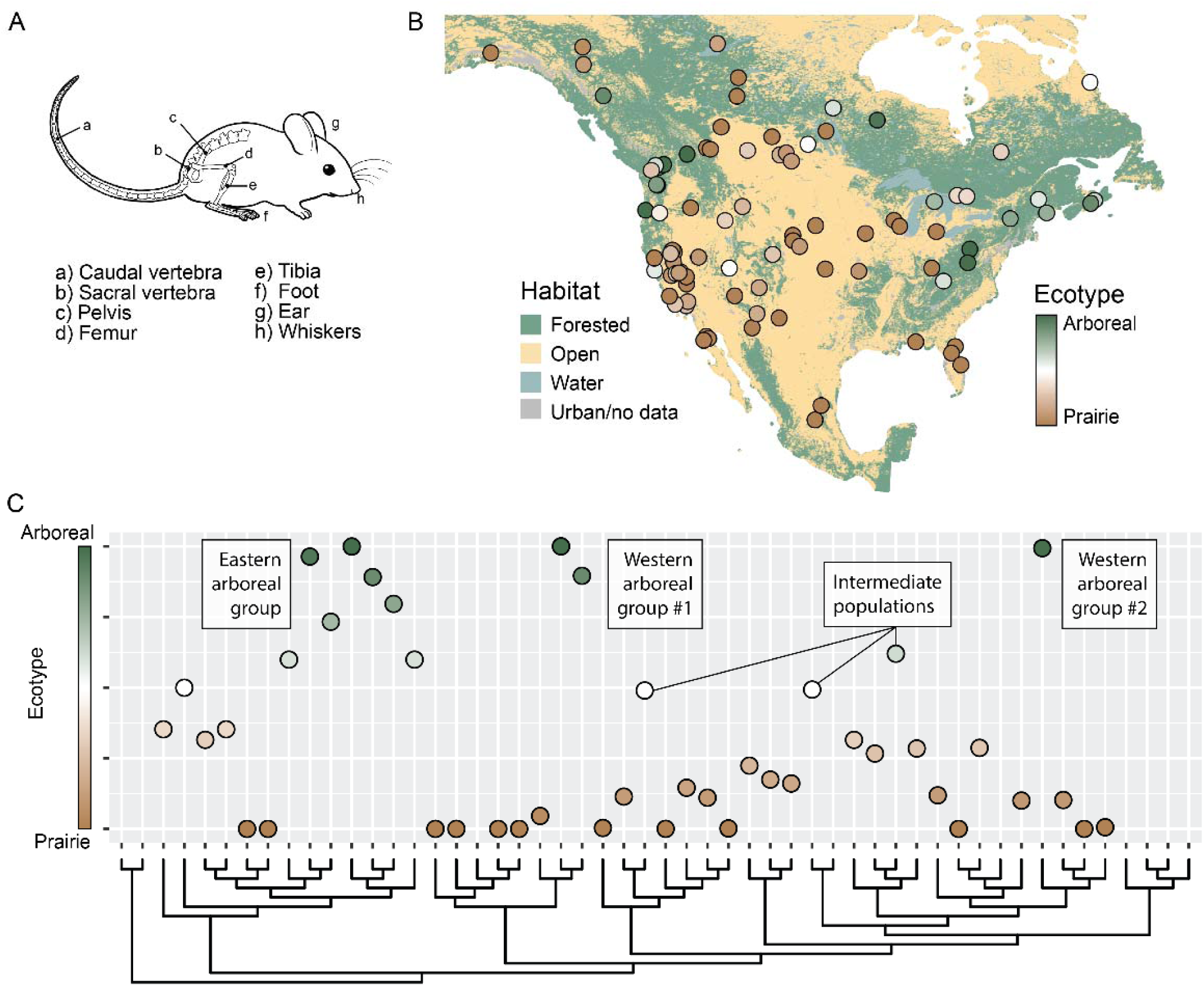
**A)** Simplified diagram of a deer mouse displaying morphological areas of interest. For brevity only general anatomical areas are annotated, not the full set of measured traits (see Table S6 for full set). **B)** Map of North America color coded by “forest” or “prairie” habitat as summarized at 3km resolution from NALCMS satellite data, overlaid with predicted ecotype of all populations with phenotypic data. **C)** Predicted ecotype an habitat mapped to SVDQuartets population tree (Table S8). Absence of predicted ecotype indicates insufficient data for prediction.

Our ecotype classification strategy suggests that the arboreal ecotype evolved at least three times independently in *P. maniculatus* (Fig. 4C). By mapping ecotype assignments onto the population-level coalescent phylogeny, we see that arboreal populations cluster into three primary groups, one eastern and two western. All of the eastern arboreal mice appear to form a single paraphyletic group, even though the most distant members of this group are separated by 2,728 km. In contrast, the two western arboreal groups are geographically close (mean pairwise distance = 732 km) yet share a deep common ancestor in our tree. The separation of the three arboreal groups in the tree suggests that the arboreal ecotype evolved independently at least three times. Alternatively, there was a single origin and repeated losses (see Discussion). We also observe several populations with partial arboreal ecotypes. In most cases this results from polymorphism, with some sampled individuals having more arboreal-like traits (e.g., longer tails) than others. Finally, prairie-like populations are more common and found at all extremes of the sampled range.

## Discussion

It is becoming increasingly common to integrate ecological and evolutionary processes, such as local adaptation, into phylogeographic studies (Hickerson *et al*. 2010; Alvarado-Serrano and Knowles 2014). Here, we draw on climate, landscape, genetic, and morphological data to show that the deer mouse has responded to past climatic shifts through widespread contraction and expansion of its range. These changes probably led to repeated phenotypic evolution upon reintroduction to similar environments. This process is demonstrated most strikingly by the repeated occurrence of arboreal and prairie ecotypes in populations that are geographically and genetically distant from each other.

*Diversity and structure in the* Peromycus maniculatus *species complex*

We recovered six highly-supported groups in our phylogenomic and population genomic analyses (Fig 2). Since the first description of *P. maniculatus* over a hundred years ago (Osgood 1909), the sheer diversity of this group has continued to challenge systematists (Bradley *et al*. 2019; Greenbaum, Honeycutt and Chirhart 2019). Our analyses show that the many morphological and regional designations of *Peromyscus* are all nested within *P. maniculatus* (Fig. 2). Some of these higher-level relationships were recovered in earlier mitochondrial phylogenies, but the precise placement of some groups - for example, whether *P. m. gambelli* is within or sister to *sonoriensis* - has changed frequently (Dragoo *et al*. 2006; Kalkvik *et al*. 2012; Natarajan *et al*. 2015; Sawyer *et al*. 2017; Bradley *et al*. 2019). Our multi-locus dataset, the largest yet for the *P. maniculatus* species complex, further modifies some of these lower-level relationships and casts doubt on previous species and subspecies designations. We also recovered significant evidence for gene flow between multiple lineages within the species complex (Fig. S6; Table S3), and laboratory crosses between distant deer mouse populations and putative species are commonplace (Bendesky *et al*. 2017; Hager *et al*. 2022; Kautt *et al*. 2024, Kinglsey *et al*. 2021). Although further sampling might unveil a working taxonomy, we propose that the evolutionary history of the *P. maniculatus* species complex will be too difficult to determine given current data and tools. We have provided an extended discussion of this in the supplementary material for the interested reader.

### Expansion and contraction through the Pleistocene

We hypothesize that rapid range expansion following the end of the Last Glacial Maximum shaped contemporary deer mouse diversity. Ecological niche models indicate that deer mice experienced reduced suitable conditions during the LGM and were probably restricted to below the 40th parallel. As the Laurentide ice sheet retreated their range began to expand (Fig. 3; Supplementary Video). Evidence from reconstructed population sizes also suggests diversification during this period (Fig. S7). The glacial-interglacial climatic oscillations of the Pleistocene have impacted many North American species in similar ways, both altering the distribution of genetic diversity and leading to range expansions and contractions (Arbogast 2007; Malaney *et al*. 2013; Jezkova *et al*. 2015; Massatti and Knowles 2016; Sim *et al*. 2016; Reid *et al*. 2019; Boria *et al*. 2021; Wooldridge *et al*. 2025). Our ENM and genetic results suggest that this scenario is true of deer mice, as has also been hypothesized in previous biogeographic work (Dragoo *et al*. 2006; Kalkvik *et al*. 2012; Sawyer *et al*. 2017; Bradley *et al*. 2019; Boria and Blois 2023). This is further corroborated by fossil evidence. Extant species of *Peromyscus*, including *P*. *maniculatus*, are common in fossil deposits dating to the late Pleistocene (∼0.129-0.012 Ma) and Holocene (∼0.012 Ma-present) (King 1968; Blois, McGuire and Hadly 2010). Success during the late Pleistocene and Holocene may have been facilitated by their broad environmental and climate tolerances, allowing *Peromyscus* to persist during the climatic oscillations of the last glacial and deglacial period (King 1968; Blois, McGuire and Hadly 2010).

### The repeated evolution of arboreal and prairie ecotypes

Range expansion and contraction during the Pleistocene glacial cycles likely led to the repeated evolution of forest and prairie ecotypes. Using both publicly available specimen records and our own phenotypic measurements of 979 specimens (Fig. 4, Fig. S10), we find pervasive, range-wide associations between forested habitat and arboreal traits, as well as the converse - associations between diverse open habitats (e.g. prairies, chaparral) and classical prairie traits. Until recently, most analyses of the arboreal and prairie ecotypes were either qualitative descriptions (King 1968), or, if quantitative, represented only select populations (Kingsley *et al*. 2017; Hager *et al*. 2022). Absent from previous work was a comprehensive, range-wide analysis of traits associated with both arboreal and prairie ecotypes, as presented here. Despite the strength and breadth of the habitat-trait associations we observe, these relationships are not perfect (Fig. 4). Mismatch between traits and the environment could be driven by any number of factors, from weak selection for arboreality to a lack of adaptive variation for selection to act upon. Alternatively, competing selection pressures may be at work, for example, selection for smaller body sizes and appendages in colder climates (i.e., Bergmann’s rule) (Guralnick *et al*. 2020). Importantly, the traits we measured only represent some features of arboreal and prairie ecotypes. While we have captured morphology that may aid navigation in complex and dense forests, it is thought that arboreal deer mice may also exhibit behavioral adaptations like skilled climbing (Horner 1954; Wecker 1963; Hager and Hoekstra 2021). The deer mouse literature is rife with evidence of heritable local adaptations (Schweizer *et al*. 2019; Linnen *et al*. 2013; Wooldridge *et al*. 2022), and it is possible that other unmeasured arboreal traits exist.

When combined with our phylogenomic analyses, this phenotypic survey suggests that the arboreal ecotype has evolved independently at least three times (Fig. 4). Ancestral state reconstruction is difficult in high gene flow and ILS systems (Revell 2025), so here we assume that the prairie ecotype represents the ancestral state based on it being more common across the landscape and in our phylogeny. During glacial cycles, deer mice potentially migrated to regions with more grassland availability (Williams 2003). Operating on this assumption, we see a single origin for the arboreal ecotype in Eastern North America, as all the arboreal populations there are closely related. In Western North America, the arboreal ecotype has probably evolved at least twice. These results add an independent arboreal lineage to the two previously identified by Kingsley et al. (2017). While it’s unlikely that further sampling will uncover arboreal lineages independent of these three, the ‘intermediate’ populations we identified could still serve as useful replicates in studies of independent trait evolution (Fig. 4). Repeatedly evolved traits are useful for understanding evolutionary genetic constraints, particularly when they occur on brief timescales between closely related lineages (Xie *et al*. 2019). However, studies of repeatedly evolved traits are often limited in the number of independent replicates (Wooldridge *et al*. 2022) or the number of distinct traits involved (Nosil *et al*. 2018). The repeated and recent evolution of multiple traits characterized here represents an ideal system for addressing these shortcomings.

## Acknowledgments

We thank M. Omura and the Harvard Museum of Comparative Zoology (MCZ) for help with accessioning specimens and providing specimen loans. We also thank the University of New Mexico Museum of Southwestern Biology (MSB), the University of Alaska Museum of the North (UAM), the Denver Museum of Nature and Science (DMNS), Natural History Museum of Los Angeles (LACM), the University of Washington Burke Museum (UWBM), and the University of California at Berkeley Museum of Vertebrate Zoology (MVZ) for loaning specimens for this research. J. Krupa, J. Orrock, J. Light, B. Danielson, R. Poulin, B. Redquest, and J. Storz all provided specimens from their personal collections. J. Gable, A. Law, P. Bhat, and S. He all contributed to whisker measurements. E. Hager helped with X-ray imaging, and both E. Hager and E. Kingsley provided several X-ray images. B. Neugeboren, K. Turner, and O. Harringmeyer helped with library preparation. O. Harringmeyer and T. Sackton provided feedback during writing of the manuscript.

TBW was supported by grants from the American Society of Mammalogists (#7623092) and the Harvard Museum of Comparative Zoology, KAG by the Program for Research in Science and Engineering (PRISE) Fellowship awarded by the Harvard College Office of Undergraduate Research and Fellowships (URAF). HEH was supported by the Howard Hughes Medical Institute. RAB was supported by an NSF Postdoctoral Research Fellowship in Biology (NSF- 2109652).

## Author Contributions

TBW identified and acquired specimens for phenotyping and genotyping. TBW, SM, CK, and KAG performed all phenotyping. TBW, RB, and AFK generated the whole genome sequence data and performed all genomic analyses. The manuscript was written by TBW, RB, and AFK with input from all authors.

## Methods

### Population sequencing and variant calling

Morphological vouchers and tissues for sequencing were provided by a series of collaborators and natural history museums (Data S1). For high coverage whole-genome sequencing (WGS), we extracted DNA from ∼20mg of liver, ear punch, or tail tip tissues. We then generated sequencing libraries from these tissues using a combination of PCR-free (KAPA HTP) and PCR-based (KAPA LTP) approaches, depending on quantity of source tissue and time of extraction. Following enzymatic fragmentation, we used size selection to enrich for a 450bp insert size and ligated Illumina adapters. We sequenced the resulting libraries using 150bp paired-end sequencing on Illumina NovaSeq S4 flowcells to achieve 15-20X coverage.

We converted raw fastq files to unmapped bam files using *FastqToSam* (*Picard* toolkit v2.27.1, 2019) and then marked Illumina adapters using *MarkIlluminaAdapters* (*Picard*). Using *SamToFastq* (*Picard*), we created interleaved fastq files and clipped adapter sequences. We mapped sequencing reads to the *P. maniculatus bairdii* reference genome (GCA_003704035.3) using *bwa-mem v0.7.17* (Li and Durbin, 2009), with –p to indicate interleaved paired-end fastq input, and –M to mark short split hits as secondary for compatibility with Picard. We then used *MergeBamAlignment* (*Picard*) to merge mapped and unmapped bam files to preserve read group information. Finally, we flagged sequencing read duplicates using *MarkDuplicates* (*Picard*), with OPTICAL_DUPLICATE_PIXEL_DISTANCE=2500 to account for artifacts generated from the patterned flowcell of the NovaSeq S4. To begin variant calling, we used *HaplotypeCaller* (GATK 4.2.6.1; Poplin et al., 2018) on the aligned bam files with the default heterozygosity prior (-hets = 0.005) and –ERC GVCF to produce per-sample gVCFs. For the X chromosome, we specified a prior input ploidy based on a comparison of coverage with the autosomes using *samtools* (v. 1.10) *depth* (Li et al., 2009). Next, we created GenomicsDB objects for each chromosome using *GenomicsDBImport* (GATK 4.2.6.1). Given the high levels of sequence polymorphism in these samples, we reduced computational time by creating GenomicsDB objects for smaller (<10MB) genomic intervals within each chromosome, which ended in natural assembly gaps, where possible. We then ran *GenotypeGVCFs* (GATK 4.2.6.1) on each subregion with “--max-alternate--alleles 4 -all-sites” to obtain raw variant calls. Subregion vcfs were then combined into whole-chromosome vcfs with *GatherVcfs* (GATK 4.2.6.1). Finally, these chromosome-level vcfs were split into indels and SNPs with *SplitVcfs* (*Picard*). We performed filtering on each variant class independently, excluding SNPs with QD < 2.0, FS > 10.0, MQ < 40.0, MQRankSum < -12.5, ReadPosRankSum < -8.0 or SOR > 3.0 and excluding INDELs with QD < 2.0, FS > 200.0, ReadPosRankSum < -20.0, SOR > 3.0. These filtering parameters were based on a combination of GATK recommendations for datasets without truth/training sets, and visual inspection of the distributions for each metric. We also set individual genotype calls to missing (./.) if the read depth at a given site was less than five using *bcftools v1.19*.

### SNP data masking

We masked regions of the genome that may be enriched for mapping and genotyping errors given our short-read sequencing approach, and known regions unlikely to be evolving under simple genetic drift. More specifically, our mask covered (i) regions of low mappability as determined with *genmap v.1.3.0* (-E 4), (ii) repetitive regions softmasked in the reference genome (determined with *WindowMasker* <u>(Morgulis et al. 2006)</u> as part of NCBI’s annotation pipeline), (iii) chromosomal inversions known to segregate in deer mice <u>(Harringmeyer and Hoekstra 2022)</u>, (iv) regions of abnormally high read depth (defined as higher than mean plus 4 standard deviations; read depth determined with *samtools depth*), (iv) exons +- 1kb in NCBI’s annotation release 102 or an in-house generated annotation <u>(Lassance et al. 2025)</u>.

### Population structure

To investigate population structure, we used two non-spatially explicit approaches and one spatially explicit method. All methods attempt to determine the number of populations within a dataset. The first non-spatial method we used was *ADMIXTURE* (v. 1.3), a maximum likelihood approach to determine the optimal number of populations (Alexander, Novembre and Lange 2009). We used 10-fold cross-validation error values across different values of K to determine the optimal number of populations. We also used the nonmodel-based method Discriminate Analysis of Principal Components (DAPC) in the *adegenet* package (Jombart 2008; Jombart and Ahmed 2011) in R (v. 4.1.1; R core team 2021) to infer structure. For the DAPC analysis, we used Bayesian information criterion to determine the optimal number of populations. Finally, we used the spatially explicit approach *TESS3r*, which incorporates geography and genotypic data, in R (Caye *et al*. 2016, 2018).

### Phylogenetic reconstructions

We first inferred phylogenetic relationships amongst individuals using two methods: 1) *IQTREE* v 2.2.2.6 (Nguyen *et al*. 2015; Minh *et al*. 2020); 2) *SVDquartets* (Chifman and Kubatko 2014). To determine if we identify mito-nuclear discordance within this species complex, we ran *IQTREE*, a software program used for large datasets that employs a maximum likelihood (ML) algorithm, on the concatenated SNPs and mitochondrial dataset (11 genes; Table S1). two datasets: 1) a ‘whole’ dataset, which consisted of all 232 ingroup + 5 outgroup individuals; and 2) a “reduced” dataset including only individuals that had higher than 70% population assignment to a single group (211 ingroup + 5 outgroup individuals) to improve the phylogenetic resolution. For the concatenated SNP dataset and the mitochondrial dataset, we used 1000 ultrafast bootstraps and the model selection tool, *ModelFinder Plus*, with an ascertainment bias correction for the SNP data (Kalyaanamoorthy *et al*. 2017; Hoang *et al*. 2018; Minh *et al*. 2020). To infer a population (‘species’) tree for this complex, we next implemented SVDquartets using a multi-species coalescent model in *PAUP\** (v4.0a) (Wilgenbusch and Swofford 2003). We utilized all quartets and 100 bootstrap replicates for both the “whole” and the “reduced” datasets. To identify gene flow between lineages, we estimated Patterson’s D statistic and f_4_-ratios using *Dsuite* (Patterson et al. 2012; Malinsky et al. 2020). We applied the Dtrios and Fbranch functions, using the Benjamini-Hochberg (BH) correction to control the false discovery rate, with a significance threshold set at p = 0.05.

### Demographic inference

We reconstructed demographic histories for representative high-coverage individuals (>19X) from the six main clades (Fig. 2) using *MSMC2* <u>(Schiffels and Wang 2020)</u>. *MSMC2* and related methods are sensitive to coverage and the effects of missing data. Therefore, after filtering for autosomal SNPs in each individual we applied three genomic masks: 1) a mask to exclude regions in the reference assembly with poor mappability (see above), 2) a mask to exclude known chromosomal inversions segregating in *P. maniculatus* (see above), and 3) a mask to exclude regions within too low (avg/2) or too high (avg*2) read depth for each individual. For this last individual-specific mask, we used the *msmc-tools* script *bamCaller.py* on the output of *bcftools mpileup -q 20 -Q 20 -C 50 -u | bcftools call -c -V indels* for each bam alignment. For each sample, we combined our input variant calls and these three masks to generate *MSMC2* input with the msmc-tools script *generate_multihetsep.py*. We created bootstrap replicates of this input data using the multihetsep_bootstrap.py, specifying -n 23 -s 5000000 –chunks_per_chromosome 10 –nr_chromosomes 20. Finally, we ran *MSMC2* on the original data and all bootstrap replicates with the time segment pattern specified as -p 25*1 + 1*2 + 1*3. All demographic histories were then visualized in R and scaled by the estimated mutation rate of 5.34 × 10^−9^ <u>(Bergeron et al. 2023)</u> and generation time of 0.5. For contextualizing these demographic histories, we retrieved marine isotope stage (MIS) geological data with the R library *gsloid* v.0.2.0 <u>(Marwick et al. 2022)</u>. We calculated Tajima’s D using a 100,000 bp sliding window for five lineages, excluding *P. melanoti*s due to sample size, within the species complex using VCFtools (v. 0.1.17; Danecek et al., 2011).

### Ecological niche modeling

We generated ecological niche models (EMNs) for *Peromyscus maniculatus* from the present to the Last Glacial Maximum (LGM [21 KYA]; at 500 year intervals) to understand how the Pleistocene glacial cycles impacted its distribution. To understand the impact of the glacial cycles on deer mice and how it could have affected diversification and range expansion/ contraction, we only included individuals labeled as *P. maniculatus* within the six mitochondrial defined clades. Occurrence localities were compiled from (Boria and Blois 2018) and this current study (Extended Data 3). To remove spatial biases, we spatially filtered the dataset to ensure no two localities were within 50 km of one another (Boria *et al*. 2014) using the R package *spThin* (Aiello-Lammens *et al*. 2015; Extended Data Table 4). For environmental data, we used Community Climate Simulation Model 3 (CCSM3; Liu et al., 2009). CCSM3 variables were downscaled to 50 x 50 km degree grid cells for North America from 21,000 years ago to the present day (Lorenz *et al*. 2016). To approximate modeling assumptions regarding dispersal and biotic interactions more closely, we delimited a custom study region by drawing a minimum convex polygon around the localities and adding a 3.0° buffer (Anderson and Raza 2010; Barve *et al*. 2011). We used a machine learning algorithm, *maxent* (v3.4.1) (Phillips, Anderson and Schapire 2006; Phillips *et al*. 2017) to infer the ENMs. We calibrated and evaluated the models using a geographically structured 5 *k-fold* approach (Radosavljevic and Anderson 2014) in the *ENMeval* package in R (v2.0.4) (Muscarella *et al*. 2014; Kass *et al*. 2023). To select species-specific model settings approximating optimal levels of complexity, we tuned model settings by varying different combinations of feature class and regularization multiplier (RM; Shcheglovitova and Anderson 2013). To identify the optimal parameter settings, we evaluated model performance using sequential criteria (minimizing overfitting and then maximizing discriminatory ability; (Shcheglovitova and Anderson 2013; Muscarella *et al*. 2014)). The optimal *Maxent* settings were Linear and Quadratic features with a RM = 2.0 and had an AUC of 0.714 and an omission rate of 0.001 (Table S5). We used the optimal settings to project the ENM into current climatic conditions and at 500-year intervals until the LGM.

### Species ranges and habitat classification

The species range for *P. maniculatus* was obtained from the IUCN Red List database (IUCN, 2021). All continental maps and bodies of water were obtained using the *ne_countries*/*ne_download* functions from the R package *rnaturalearth* v.0.1.0 (South 2017). Continent-wide habitat classifications were performed by summarizing land cover types from the North American Land Change Monitoring System (NALCMS, 2010). Based on NALCMS habitat descriptions, we considered six habitat classes with predominantly tall (>3m) and dense vegetation to be “forest” (temperate or sub-polar needleleaf forest, sub-polar taiga needleleaf forest, tropical or sub-tropical broadleaf evergreen forest, tropical or sub-tropical deciduous forest, temperate or sub-polar broadleaf deciduous forest, mixed forest), eleven habitat classes with predominantly short (<3m) or absent vegetation as “prairie” (tropical or sub-tropical shrubland, temperate or sub-polar shrubland, tropical or sub-tropical grassland, temperate or sub-polar grassland, sub-polar or polar shrubland-lichen-moss, sub-polar or polar grassland-lichen-moss, sub-polar or polar barren-lichen-moss, wetland, cropland, barren lands), and three habitat classes as “NA” (urban and built-up, water, snow and ice). For lower-memory computation, we transformed the NALCMS resolution from 30m to 3km (100-fold reduction) by reporting the most common habitat class (forest, prairie, NA) in each 3km zone using the function *aggregate* from the package *terra* v.3.4.5 (Hijmans 2023)

### Measurement of ecotype traits

We obtained three primary types of phenotypic data from each specimen, where possible. First, we obtained external measurements taken upon initial trapping or accession of the specimen, namely total length (mm), body length (mm), tail length (mm), ear length (mm), hind foot length (mm), weight (g), and sex. These measurements were either obtained directly from the collector, or from *P. maniculatus* museum records available via the Arctos (https://arctosdb.org/) or Vertnet (http://vertnet.org/) databases. Second, skeletal measurements were obtained from X-ray imaging of whole-body specimens. Imaging was performed using a KEVEK X-ray source and Varian digital imaging panel (Varian Medical Systems, Inc.), and subsequent image analyses were performed in FIJI (Schindelin *et al*. 2012). Third, whisker measurements were obtained from both whole-body specimens and prepared flatskins. Where possible, the lengths of the alpha, beta, gamma, and delta caudal offset whiskers were recorded from both left and right flanks of the specimen, following protocols described in Gable (2020). A list of phenotypes we directly measured can be found in Table S6. Full trait data for both the museum and detailed trait datasets will be available on Dryad upon publication (10.5061/dryad.612jm64jf).

Prior to downstream analyses, we filtered out implausible values likely caused by measurement error. Specifically, we removed individuals with a total body length less than 115 mm, as such individuals were likely to be juveniles. If fewer than 20 caudal vertebrae could be measured, we considered observations pertaining to the total caudal vertebrae number and terminal caudal vertebrae lengths to be incomplete. For all remaining measurements, we removed outliers by discarding values outside the 1^st^ and 3^rd^ quartiles -/+ two times the interquartile range, assuming that these extreme observations were most likely to indicate measurement error, a common feature of museum records. Finally, we excluded any traits for which less than 25% of the individuals had observations, with the exception of terminal caudal vertebrae for which the absence of any value was informative.

### Linear discriminant analysis and phenotypic PCA

#### Imputation of missing values

To allow downstream phenotypic comparisons and analyses, we first had to account for missing observations in our phenotypic dataset. Using the function *amelia* from the R package *Amelia* v.1.8.0 (Honaker, King and Blackwell), we imputed missing trait values for each individual by computing means from 100 independent imputations on the core dataset. Imputation quality was assessed by manually inspecting distributions of observed and imputed values and ensuring that the distributions were approximately matched.

#### Linear discriminant analysis (LDA) and ecotype classification

In order to determine whether a population’s morphology more closely resembled forest or prairie ecotypes, we constructed a linear discriminant model based on a core set of well-studied representatives for each ecotype. More specifically, we used one western forest-prairie pair (Hager and Hoekstra 2021) and one eastern pair (Kingsley *et al*. 2021). Prior to model fitting, we centered each trait variable to allow easier downstream comparison of variable contributions. We then fit the model using the *lda* function from the R package *MASS* v.7.3.53.1 (Venables and Ripley 2002), setting ‘arboreal’ and ‘prairie’ as the response classification variable, and all phenotypic observations as the predictors. After performing leave-one-out-cross-validation with the training dataset, we observed strong model accuracy (90.8%), and proceeded with ecotype prediction (Fig. S10).

Posterior ecotype probabilities were predicted for each individual using *predict(., type = “posterior”)* from the base R package *stats* v.4.0.5, assuming prior probabilities for ‘forest’ and ‘prairie’ based on proportions individual observed in the training data. We then determined ecotype probabilities for each population by calculating the mean across all individuals in each population. This mean ecotype posterior probability was then used for all downstream phenotypic analyses, with values of ‘0’ and ‘1’ representing prairie and forest, respectively.

#### Phenotypic PCA

To further explore trait variation among populations, we performed principal components analysis (PCA) using the *PCA* function from the R package *FactoMineR* v.2.4 (Lé, Josse and Husson). PCA was performed on individual samples, but visualized at the population level (Fig. 4A) using the center-of-gravity calculations returned by *FactoMineR::PCA*.

## Supplementary Materials

### Taxonomic implications of the Peromycus species complex

Since the first description of *P. maniculatus* over a hundred years ago (Osgood 1909), the sheer diversity within the complex has continued to challenge systematists’ attempts to delineate species within it. (Bradley *et al*. 2019; Greenbaum, Honeycutt and Chirhart 2019). Here, by applying a suite of phylogenomic and population genomic analyses to the largest multi-locus dataset to date of the *P. maniculatus* species complex, we consistently recovered six distinct clades of deer mice (Figs 1 & 2). Several of these clades have frequently been labelled as the distinct species *P. keeni*, *P. labecula, P. polionotus*, and *P. sonoriensis*, yet we see that they nest within *P. maniculatus* based on autosomal and mitochondrial evidence (Fig. 2). These results represent whole-genome confirmation of patterns reported from previous mitochondrial phylogenies (Dragoo *et al*. 2006; Kalkvik *et al*. 2012; Natarajan *et al*. 2015; Sawyer *et al*. 2017; Bradley *et al*. 2019). One clade we report represents a previously undescribed lineage, *P. m.* Midwest. This group was identified by every population genetic analysis (Fig. S1), was sister to all other clades in the mitochondrial tree (Fig, 2) and formed a monophyletic clade in the nuclear phylogenetic trees (Fig. 2; Fig. S4; Fig. S5).

However, several notable differences between the mitochondrial and nuclear trees shed light on previous work in *P. maniculatus* systematics. Autosomal evidence indicates *P. m. melanotis* is sister to all other deermouse clades, whereas the mitochondrial tree shows it nested within *P. maniculatus* (Fig. 2; Fig. S4; Fig. S5). Unsurprisingly, the placement of *P. m. melanotis* has changed over time, with some studies indicating it was nested within the deer mouse species complex as in our mitochondrial tree (Fig. 2) (Dragoo *et al*.2006; Natarajan *et al*. 2015) whereas other mitochondrial studies showed it was sister to all other deermouse clades - the nuclear relationship we observe (Kalkvik *et al*. 2012; Bradley *et al*. 2019; Greenbaum, Honeycutt and Chirhart 2019). Similarly, *Peromyscus m. labecula* clustered with *P. m. sonoriensis* in all population genomic analyses and in the mitochondrial tree, but the ML trees placed these individuals as sister to all groups after *P. m. melanotis* and the population tree put this group sister to the eastern and western clades (Fig. 2; Fig. S4; Fig. S5).

At times our mitochondrial and nuclear analyses were in agreement yet differed from previous findings. The primary example of this is the ‘oldfield mouse’ *P. m. polionotus* (or *P. polionotus*), which inhabits the southeastern United States where traditionally recognized deer mouse populations are absent. Although previous mitochondrial work indicated this group was sister to western *P. maniculatus* clades, both mitochondrial and nuclear trees place this lineage as sister to *P. m. maniculatus* (the northeastern clade; Fig. 2; Fig. S4; Fig. S5). Additionally, recent taxonomic revisions elevated *P. m. gambelli* (a southern California mitochondrial clade) to the species level (Greenbaum *et al*. 2017; Bradley *et al*. 2019); however, all population genetic and phylogenetic analyses indicate *P. m. gambelli* is nested within *P. m. sonoriensis*, corroborating recent findings (Boria and Blois 2023). Finally, in northwestern North America three *Peromyscus* species have been described: *arcticus*, *keeni*, and the aforementioned *sonoriensis (Sawyer et al. 2017)*. Previous mitochondrial trees show *arcticus* and *keeni* as sister to *P. m. gambelli* (Bradley *et al*. 2019), while more recent analyses based on nuclear data suggest these clades are sister to *P. m. sonoriensis* (Sawyer *et al*. 2017; Boria and Blois 2023). The nesting of *P. m. gambelli* within *P. m. sonoriensis* we describe above may explain these alternate placements. Nevertheless, here we show that *arcticus* and *keeni* cluster together in population genetic analyses and form a monophyletic clade nested within the *P. maniculatus* complex in nuclear trees (Fig. 2, Fig. S4; Fig. S5). The mitochondrial tree disrupts this monophyly, but still shows *keeni* and *arcticus* as closely related to each other and to some individuals of *sonoriensis* (Fig. 2). Additional work may determine if *arcticus*, *keeni*, and these individuals of *sonoriensis* represent unique species or form one clade.

Hybridization between recently diverged species happens readily in nature (Payseur and Rieseberg 2016), and we recovered significant gene flow between multiple lineages within the species complex (Fig. S6; Table S5). Further, laboratory experiments have shown that *P. polionotus* can be crossed with other deer mouse populations and produce viable offspring (Bendesky *et al*. 2017; Kautt *et al*. 2024). Indeed, laboratory crosses between deer mouse populations are commonplace (Hager *et al*. 2022), even those that are geographically or genetically distant (Kingsley *et al*. 2021), and in our own analyses we identified several clearly admixed individuals (Fig. S2; Table S4). Post-copulation speciation mechanisms have begun to evolve over time between members of the species group. For example, only a female *P. maniculatus* and male *P. polionotus* produce viable offspring (Moore 1965; Vrana *et al*. 2014). Sperm competition between individuals from *P. polionotus* and *P. maniculatus* indicate that conspecific sperm aggregate allowing for discrimination between species and *P. maniculatus* (a promiscuous species) showed sperm aggregation between individuals post-copulation (Fisher and Hoekstra 2010). Although the biological species concept is only one potential criterion to delimit species, these phenomena - in light of our own genomic results - suggest that delineating all species in the *Peromyscus maniculatus* complex may prove intractable. It is likely that we are witnessing groups in the early stages of speciation, in which case many species concepts will inevitably fall short.

---

**Table S1.** Mitochondrial partition file.

| Molecule | Gene | Start | End |
| --- | --- | --- | --- |
| DNA | ND4L | 9925 | 10221 |
| DNA | ND1 | 2765 | 3719 |
| DNA | ND2 | 3930 | 4967 |
| DNA | COX1 | 5348 | 6892 |
| DNA | COX2 | 7031 | 7714 |
| DNA | ATP6 | 7946 | 8628 |
| DNA | COX3 | 8628 | 9411 |
| DNA | ND3 | 9481 | 9828 |
| DNA | ND4 | 10204 | 11583 |
| DNA | ND5 | 11781 | 13600 |
| DNA | CYTB | 14199 | 15342 |

**Table S2.** The BIC results from DAPC on *Peromyscus maniculatus* across its range in North America.

| Populations | BIC |
| --- | --- |
| K=1 | 1833.223 |
| K=2 | 1809.532 |
| K=3 | 1805.218 |
| K=4 | 1798.578 |
| K=5 | 1793.254 |
| K=6 | 1792.456 |
| K=7 | 1789.561 |
| K=8 | 1789.72 |
| K=9 | 1790.543 |
| K=10 | 1791.036 |

**Table S3.**
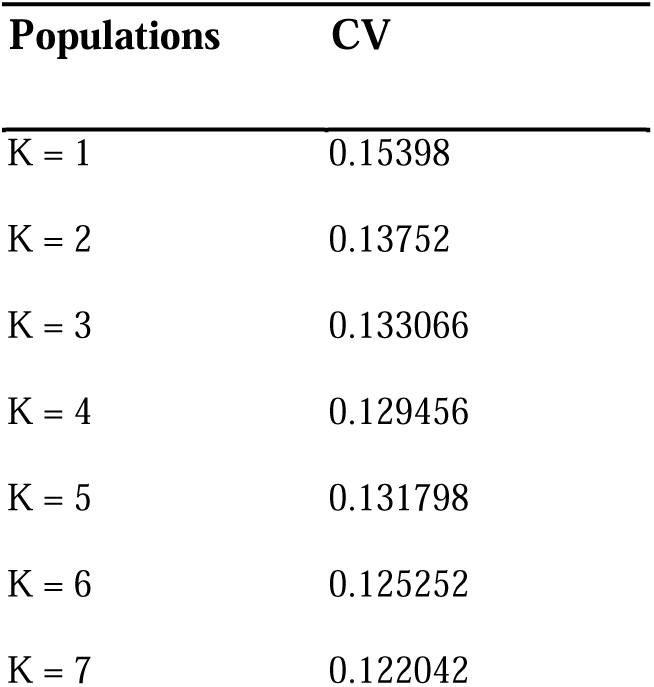
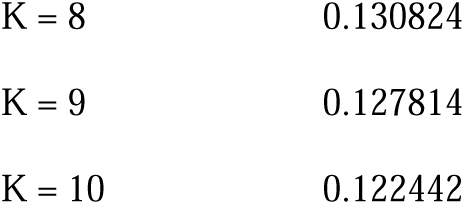
The 10-fold cross validation results from ADMIXTURE on *Peromyscus maniculatus* across its range in North America.

**Table S4.** The ancestry fractions from ADMIXTURE, K = 7, for *Peromyscus maniculatus* across its range in North America.

| sampleID | K1 | K2 | K3 | K4 | K5 | K6 | K7 |
| --- | --- | --- | --- | --- | --- | --- | --- |
| 4434 | 0.00001 | 0.99994 | 0.00001 | 0.00001 | 0.00001 | 0.00001 | 0.00001 |
| 7817 | 0.00001 | 0.99994 | 0.00001 | 0.00001 | 0.00001 | 0.00001 | 0.00001 |
| 68144 | 0.00001 | 0.99994 | 0.00001 | 0.00001 | 0.00001 | 0.00001 | 0.00001 |
| 68146 | 0.00001 | 0.999938 | 0.00001 | 0.00001 | 0.00001 | 0.000012 | 0.00001 |
| 68147 | 0.00001 | 0.99994 | 0.00001 | 0.00001 | 0.00001 | 0.00001 | 0.00001 |
| 68148 | 0.00001 | 0.99994 | 0.00001 | 0.00001 | 0.00001 | 0.00001 | 0.00001 |
| 68149 | 0.00001 | 0.99994 | 0.00001 | 0.00001 | 0.00001 | 0.00001 | 0.00001 |
| 69677 | 0.00001 | 0.99994 | 0.00001 | 0.00001 | 0.00001 | 0.00001 | 0.00001 |
| 69679 | 0.00001 | 0.99994 | 0.00001 | 0.00001 | 0.00001 | 0.00001 | 0.00001 |
| 69686 | 0.00001 | 0.99994 | 0.00001 | 0.00001 | 0.00001 | 0.00001 | 0.00001 |
| 77713 | 0.989862 | 0.00001 | 0.00001 | 0.00001 | 0.010088 | 0.00001 | 0.00001 |
| 77716 | 0.588949 | 0.00001 | 0.376749 | 0.00001 | 0.034262 | 0.00001 | 0.00001 |
| 77725 | 0.781221 | 0.00001 | 0.00001 | 0.00001 | 0.218729 | 0.00001 | 0.00001 |
| 78153 | 0.745104 | 0.00001 | 0.00001 | 0.00001 | 0.254846 | 0.00001 | 0.00001 |
| 78154 | 0.748949 | 0.00001 | 0.00001 | 0.00001 | 0.251001 | 0.00001 | 0.00001 |
| 78155 | 0.72001 | 0.00001 | 0.00001 | 0.00001 | 0.27994 | 0.00001 | 0.00001 |
| 04_SW_F_ERH21 | 0.00001 | 0.00001 | 0.00001 | 0.00001 | 0.00001 | 0.99994 | 0.00001 |
| 05_SW_M_ERH27 | 0.99994 | 0.00001 | 0.00001 | 0.00001 | 0.00001 | 0.00001 | 0.00001 |
| 06_SW_M_ERH36 | 0.95273 | 0.00001 | 0.00001 | 0.00001 | 0.04722 | 0.00001 | 0.00001 |
| 07_BK_M_ERH108 | 0.99994 | 0.00001 | 0.00001 | 0.00001 | 0.00001 | 0.00001 | 0.00001 |
| 08_SW_M_ERH52 | 0.99994 | 0.00001 | 0.00001 | 0.00001 | 0.00001 | 0.00001 | 0.00001 |
| 10_SW_M_ERH19 | 0.99994 | 0.00001 | 0.00001 | 0.00001 | 0.00001 | 0.00001 | 0.00001 |
| 18_SW_F_ERH48 | 0.00001 | 0.00001 | 0.00001 | 0.00001 | 0.00001 | 0.99994 | 0.00001 |
| 19_SW_M_ERH26 | 0.976023 | 0.00001 | 0.00001 | 0.00001 | 0.023927 | 0.00001 | 0.00001 |
| 20_SW_F_ERH14 | 0.00001 | 0.00001 | 0.00001 | 0.00001 | 0.00001 | 0.99994 | 0.00001 |
| 22_SW_M_ERH25 | 0.00001 | 0.00001 | 0.00001 | 0.00001 | 0.00001 | 0.99994 | 0.00001 |
| 27_BK_F_ERH87 | 0.00001 | 0.00001 | 0.00001 | 0.00001 | 0.00001 | 0.99994 | 0.00001 |
| 28_BK_F_ERH99 | 0.00001 | 0.00001 | 0.00001 | 0.00001 | 0.00001 | 0.99994 | 0.00001 |
| 29_BK_M_ERH95 | 0.00001 | 0.00001 | 0.00001 | 0.00001 | 0.00001 | 0.99994 | 0.00001 |
| 31_SW_F_ERH38 | 0.99994 | 0.00001 | 0.00001 | 0.00001 | 0.00001 | 0.00001 | 0.00001 |
| 34_SW_F_ERH42 | 0.978284 | 0.00001 | 0.00001 | 0.00001 | 0.021666 | 0.00001 | 0.00001 |
| 35_BK_M_ERH97 | 0.987184 | 0.00001 | 0.00001 | 0.00001 | 0.012766 | 0.00001 | 0.00001 |
| 36_BK_M_ERH100 | 0.983617 | 0.00001 | 0.00001 | 0.00001 | 0.016333 | 0.00001 | 0.00001 |
| 37_BK_M_ERH110 | 0.00001 | 0.00001 | 0.00001 | 0.00001 | 0.00001 | 0.99994 | 0.00001 |
| 38_SW_F_ERH12 | 0.00001 | 0.000012 | 0.00001 | 0.00001 | 0.00001 | 0.999938 | 0.00001 |
| 39_BK_F_ERH83 | 0.00001 | 0.00001 | 0.00001 | 0.00001 | 0.00001 | 0.99994 | 0.00001 |
| 40_BK_F_ERH88 | 0.00001 | 0.00001 | 0.00001 | 0.00001 | 0.00001 | 0.99994 | 0.00001 |
| 41_BK_M_ERH89 | 0.99994 | 0.00001 | 0.00001 | 0.00001 | 0.00001 | 0.00001 | 0.00001 |
| 42_BK_F_ERH90 | 0.999936 | 0.00001 | 0.00001 | 0.00001 | 0.000014 | 0.00001 | 0.00001 |
| 43_BK_F_ERH96 | 0.99994 | 0.00001 | 0.00001 | 0.00001 | 0.00001 | 0.00001 | 0.00001 |
| 44_BK_F_ERH107 | 0.00001 | 0.00001 | 0.00001 | 0.00001 | 0.00001 | 0.99994 | 0.00001 |
| 476_3_1 | 0.99994 | 0.00001 | 0.00001 | 0.00001 | 0.00001 | 0.00001 | 0.00001 |
| 477_1_4 | 0.953276 | 0.00001 | 0.00001 | 0.00001 | 0.046674 | 0.00001 | 0.00001 |
| 479_2_1 | 0.99994 | 0.00001 | 0.00001 | 0.00001 | 0.00001 | 0.00001 | 0.00001 |
| 48_SW_M_ERH43 | 0.99994 | 0.00001 | 0.00001 | 0.00001 | 0.00001 | 0.00001 | 0.00001 |
| BD_A | 0.99994 | 0.00001 | 0.00001 | 0.00001 | 0.00001 | 0.00001 | 0.00001 |
| BD_C | 0.99994 | 0.00001 | 0.00001 | 0.00001 | 0.00001 | 0.00001 | 0.00001 |
| BD_D | 0.99994 | 0.00001 | 0.00001 | 0.00001 | 0.00001 | 0.00001 | 0.00001 |
| DMDM01 | 0.99994 | 0.00001 | 0.00001 | 0.00001 | 0.00001 | 0.00001 | 0.00001 |
| DMNS_13043 | 0.99994 | 0.00001 | 0.00001 | 0.00001 | 0.00001 | 0.00001 | 0.00001 |
| DMNS_13046 | 0.99994 | 0.00001 | 0.00001 | 0.00001 | 0.00001 | 0.00001 | 0.00001 |
| DMNS_13056 | 0.99994 | 0.00001 | 0.00001 | 0.00001 | 0.00001 | 0.00001 | 0.00001 |
| DMNS_14161 | 0.99994 | 0.00001 | 0.00001 | 0.00001 | 0.00001 | 0.00001 | 0.00001 |
| DMNS_14162 | 0.00001 | 0.00001 | 0.00001 | 0.00001 | 0.00001 | 0.99994 | 0.00001 |
| DMNS_17655 | 0.730847 | 0.00001 | 0.00001 | 0.00001 | 0.269103 | 0.00001 | 0.00001 |
| DMWB01 | 0.730075 | 0.00001 | 0.00001 | 0.00001 | 0.269875 | 0.00001 | 0.00001 |
| DMWB03 | 0.748452 | 0.00001 | 0.00001 | 0.00001 | 0.251498 | 0.00001 | 0.00001 |
| ERH46 | 0.99994 | 0.00001 | 0.00001 | 0.00001 | 0.00001 | 0.00001 | 0.00001 |
| ERH53 | 0.00001 | 0.000012 | 0.00001 | 0.00001 | 0.00001 | 0.999938 | 0.00001 |
| ERH93 | 0.441239 | 0.00001 | 0.014349 | 0.544372 | 0.00001 | 0.00001 | 0.00001 |
| ERH94 | 0.435296 | 0.00001 | 0.004582 | 0.560082 | 0.00001 | 0.00001 | 0.00001 |
| FSB11 | 0.446405 | 0.00001 | 0.019289 | 0.534267 | 0.00001 | 0.00001 | 0.00001 |
| FSB25 | 0.00001 | 0.00001 | 0.00001 | 0.00001 | 0.99994 | 0.00001 | 0.00001 |
| FSB37 | 0.678865 | 0.00001 | 0.067208 | 0.015805 | 0.00001 | 0.00001 | 0.238091 |
| FSB42 | 0.721864 | 0.00001 | 0.00001 | 0.004758 | 0.273339 | 0.00001 | 0.00001 |
| FSB43 | 0.374749 | 0.00001 | 0.00001 | 0.00001 | 0.625201 | 0.00001 | 0.00001 |
| FSB48 | 0.99994 | 0.00001 | 0.00001 | 0.00001 | 0.00001 | 0.00001 | 0.00001 |
| FSB51 | 0.971646 | 0.00001 | 0.00001 | 0.028304 | 0.00001 | 0.00001 | 0.00001 |
| FSB52 | 0.323109 | 0.00001 | 0.00001 | 0.00257 | 0.674282 | 0.00001 | 0.00001 |
| FSB53 | 0.465336 | 0.00001 | 0.00001 | 0.013538 | 0.521086 | 0.00001 | 0.00001 |
| FSB6 | 0.744168 | 0.00001 | 0.033615 | 0.014928 | 0.00001 | 0.00001 | 0.207259 |
| FSB64 | 0.7159 | 0.00001 | 0.034953 | 0.030809 | 0.00001 | 0.00001 | 0.218307 |
| FSB66 | 0.00001 | 0.00001 | 0.99994 | 0.00001 | 0.00001 | 0.00001 | 0.00001 |
| FSB67 | 0.00001 | 0.00001 | 0.99994 | 0.00001 | 0.00001 | 0.00001 | 0.00001 |
| FSB7 | 0.00001 | 0.00001 | 0.99994 | 0.00001 | 0.00001 | 0.00001 | 0.00001 |
| HV01 | 0.00001 | 0.00001 | 0.99994 | 0.00001 | 0.00001 | 0.00001 | 0.00001 |
| HV02 | 0.00001 | 0.00001 | 0.99994 | 0.00001 | 0.00001 | 0.00001 | 0.00001 |
| HV03 | 0.00001 | 0.00001 | 0.99994 | 0.00001 | 0.00001 | 0.00001 | 0.00001 |
| KMT04 | 0.00001 | 0.00001 | 0.99994 | 0.00001 | 0.00001 | 0.00001 | 0.00001 |
| KMT05 | 0.00001 | 0.00001 | 0.99994 | 0.00001 | 0.00001 | 0.00001 | 0.00001 |
| KMT13 | 0.00001 | 0.00001 | 0.99994 | 0.00001 | 0.00001 | 0.00001 | 0.00001 |
| KMT14 | 0.00001 | 0.00001 | 0.99994 | 0.00001 | 0.00001 | 0.00001 | 0.00001 |
| KMT19 | 0.00001 | 0.00001 | 0.99994 | 0.00001 | 0.00001 | 0.00001 | 0.00001 |
| KMT62 | 0.00001 | 0.00001 | 0.99994 | 0.00001 | 0.00001 | 0.00001 | 0.00001 |
| KMT63 | 0.00001 | 0.00001 | 0.99994 | 0.00001 | 0.00001 | 0.00001 | 0.00001 |
| KMT68 | 0.00001 | 0.00001 | 0.99994 | 0.00001 | 0.00001 | 0.00001 | 0.00001 |
| KMT69 | 0.00001 | 0.00001 | 0.99994 | 0.00001 | 0.00001 | 0.00001 | 0.00001 |
| KMT70 | 0.979179 | 0.00001 | 0.00001 | 0.000018 | 0.020763 | 0.00001 | 0.00001 |
| LFR1 | 0.980926 | 0.00001 | 0.00001 | 0.00001 | 0.019024 | 0.00001 | 0.00001 |
| LFR2 | 0.99994 | 0.00001 | 0.00001 | 0.00001 | 0.00001 | 0.00001 | 0.00001 |
| LFR3 | 0.872435 | 0.00001 | 0.00001 | 0.127515 | 0.00001 | 0.00001 | 0.00001 |
| Mamm_66692 | 0.00001 | 0.00001 | 0.00001 | 0.00001 | 0.99994 | 0.00001 | 0.00001 |
| Mamm_66712 | 0.00001 | 0.00001 | 0.00001 | 0.00001 | 0.99994 | 0.00001 | 0.00001 |
| Mamm_66713 | 0.00002 | 0.00001 | 0.00001 | 0.00001 | 0.99993 | 0.00001 | 0.00001 |
| MSB_Mamm_156126 | 0.00001 | 0.00001 | 0.00001 | 0.00001 | 0.99994 | 0.00001 | 0.00001 |
| MSB_Mamm_156153 | 0.00001 | 0.00001 | 0.00001 | 0.00001 | 0.99994 | 0.00001 | 0.00001 |
| MSB_Mamm_192746 | 0.00001 | 0.00001 | 0.00001 | 0.00001 | 0.99994 | 0.00001 | 0.00001 |
| MSB_Mamm_192747 | 0.00001 | 0.00001 | 0.00001 | 0.00001 | 0.99994 | 0.00001 | 0.00001 |
| MSB_Mamm_192748 | 0.00001 | 0.00001 | 0.00001 | 0.00001 | 0.99994 | 0.00001 | 0.00001 |
| MSB_Mamm_228874 | 0.00001 | 0.00001 | 0.00001 | 0.00001 | 0.99994 | 0.00001 | 0.00001 |
| MSB_Mamm_228891 | 0.00001 | 0.00001 | 0.00001 | 0.00001 | 0.99994 | 0.00001 | 0.00001 |
| MSB_Mamm_229272 | 0.00001 | 0.00001 | 0.00001 | 0.00001 | 0.99994 | 0.00001 | 0.00001 |
| MSB_Mamm_240535 | 0.00001 | 0.00001 | 0.00001 | 0.00001 | 0.99994 | 0.00001 | 0.00001 |
| MSB_Mamm_240538 | 0.00001 | 0.00001 | 0.00001 | 0.00001 | 0.99994 | 0.00001 | 0.00001 |
| MSB_Mamm_240545 | 0.00001 | 0.00001 | 0.00001 | 0.00001 | 0.99994 | 0.00001 | 0.00001 |
| MSB_Mamm_280721 | 0.00001 | 0.00001 | 0.00001 | 0.00001 | 0.99994 | 0.00001 | 0.00001 |
| MSB_Mamm_280893 | 0.99994 | 0.00001 | 0.00001 | 0.00001 | 0.00001 | 0.00001 | 0.00001 |
| MSB_Mamm_280973 | 0.99994 | 0.00001 | 0.00001 | 0.00001 | 0.00001 | 0.00001 | 0.00001 |
| MSB_Mamm_295115 | 0.99994 | 0.00001 | 0.00001 | 0.00001 | 0.00001 | 0.00001 | 0.00001 |
| MSB_Mamm_295124 | 0.00001 | 0.00001 | 0.00001 | 0.99994 | 0.00001 | 0.00001 | 0.00001 |
| MSB_Mamm_295253 | 0.00001 | 0.00001 | 0.00001 | 0.99994 | 0.00001 | 0.00001 | 0.00001 |
| MSB_Mamm_72249 | 0.00001 | 0.00001 | 0.00001 | 0.99994 | 0.00001 | 0.00001 | 0.00001 |
| MSB_Mamm_72255 | 0.00001 | 0.00001 | 0.00001 | 0.00001 | 0.00001 | 0.99994 | 0.00001 |
| MSB_Mamm_73957 | 0.00001 | 0.00001 | 0.00001 | 0.00001 | 0.00001 | 0.99994 | 0.00001 |
| MSB_Mamm_73971 | 0.00001 | 0.008785 | 0.00001 | 0.00001 | 0.00001 | 0.991165 | 0.00001 |
| MSB_Mamm_76796 | 0.99994 | 0.00001 | 0.00001 | 0.00001 | 0.00001 | 0.00001 | 0.00001 |
| MSB_Mamm_76804 | 0.868385 | 0.00001 | 0.00001 | 0.00001 | 0.00001 | 0.00001 | 0.131565 |
| MSB_Mamm_76807 | 0.874724 | 0.00001 | 0.00001 | 0.003198 | 0.00001 | 0.00001 | 0.122038 |
| MSB_Mamm_81004 | 0.99994 | 0.00001 | 0.00001 | 0.00001 | 0.00001 | 0.00001 | 0.00001 |
| MSB_Mamm_90207 | 0.99994 | 0.00001 | 0.00001 | 0.00001 | 0.00001 | 0.00001 | 0.00001 |
| MSB_Mamm_90211 | 0.99994 | 0.00001 | 0.00001 | 0.00001 | 0.00001 | 0.00001 | 0.00001 |
| MSB_Mamm_90220 | 0.99994 | 0.00001 | 0.00001 | 0.00001 | 0.00001 | 0.00001 | 0.00001 |
| MSB_Mamm_273971 | 0.99994 | 0.00001 | 0.00001 | 0.00001 | 0.00001 | 0.00001 | 0.00001 |
| MSB_Mamm_273972 | 0.99994 | 0.00001 | 0.00001 | 0.00001 | 0.00001 | 0.00001 | 0.00001 |
| MSB_Mamm_274169 | 0.99994 | 0.00001 | 0.00001 | 0.00001 | 0.00001 | 0.00001 | 0.00001 |
| MSB_Mamm_274202 | 0.99994 | 0.00001 | 0.00001 | 0.00001 | 0.00001 | 0.00001 | 0.00001 |
| MSB_Mamm_280706 | 0.99994 | 0.00001 | 0.00001 | 0.00001 | 0.00001 | 0.00001 | 0.00001 |
| MSB_Mamm_280708 | 0.99994 | 0.00001 | 0.00001 | 0.00001 | 0.00001 | 0.00001 | 0.00001 |
| MSB_Mamm_280720 | 0.659106 | 0.00001 | 0.00001 | 0.340844 | 0.00001 | 0.00001 | 0.00001 |
| MSB_Mamm_280729 | 0.686403 | 0.00001 | 0.00001 | 0.313547 | 0.00001 | 0.00001 | 0.00001 |
| MSB_Mamm_280948 | 0.653782 | 0.00001 | 0.006938 | 0.339239 | 0.000011 | 0.00001 | 0.00001 |
| MSB_Mamm_280961 | 0.795339 | 0.00001 | 0.00001 | 0.00001 | 0.00001 | 0.00001 | 0.204611 |
| MSB_Mamm_280963 | 0.79674 | 0.00001 | 0.00001 | 0.00001 | 0.00001 | 0.00001 | 0.20321 |
| MSB_Mamm_280964 | 0.00001 | 0.00001 | 0.00001 | 0.025069 | 0.974881 | 0.00001 | 0.00001 |
| MSB_Mamm_280968 | 0.00001 | 0.00001 | 0.00001 | 0.00001 | 0.00001 | 0.00001 | 0.99994 |
| MSB_Mamm_280977 | 0.00001 | 0.00001 | 0.00001 | 0.00001 | 0.00001 | 0.00001 | 0.99994 |
| sbm_31 | 0.00001 | 0.00001 | 0.00001 | 0.00001 | 0.00001 | 0.00001 | 0.99994 |
| UMNH_34860 | 0.237584 | 0.00001 | 0.00001 | 0.00001 | 0.00001 | 0.762366 | 0.00001 |
| UMNH_34865 | 0.076309 | 0.00001 | 0.923641 | 0.00001 | 0.00001 | 0.00001 | 0.00001 |
| UMNH_34885 | 0.063551 | 0.00001 | 0.936399 | 0.00001 | 0.00001 | 0.00001 | 0.00001 |
| MSB_Mamm_47768 | 0.060365 | 0.00001 | 0.939585 | 0.00001 | 0.00001 | 0.00001 | 0.00001 |
| MSB_Mamm_47769 | 0.501672 | 0.00001 | 0.187777 | 0.00001 | 0.00001 | 0.00001 | 0.310511 |
| MSB_Mamm_47770 | 0.582463 | 0.00001 | 0.112499 | 0.00001 | 0.00001 | 0.00001 | 0.304997 |
| MSB_Mamm_86178 | 0.027751 | 0.00001 | 0.124022 | 0.00001 | 0.00001 | 0.848187 | 0.00001 |
| MSB_Mamm_86179 | 0.078862 | 0.00001 | 0.185889 | 0.00001 | 0.00001 | 0.735209 | 0.00001 |
| MSB_Mamm_86180 | 0.066065 | 0.00001 | 0.189503 | 0.00001 | 0.00001 | 0.744392 | 0.00001 |
| MSB_Mamm_91413 | 0.499504 | 0.00001 | 0.028108 | 0.025667 | 0.00001 | 0.349223 | 0.097478 |
| MSB_Mamm_91414 | 0.085175 | 0.00001 | 0.00001 | 0.00001 | 0.00001 | 0.914775 | 0.00001 |
| MSB_Mamm_91415 | 0.223824 | 0.00001 | 0.00001 | 0.00001 | 0.00001 | 0.776126 | 0.00001 |
| 02_NB_F_EPK14 | 0.101159 | 0.00001 | 0.023255 | 0.00001 | 0.00001 | 0.875546 | 0.00001 |
| 09_NB_M_EPK07 | 0.095823 | 0.00001 | 0.003928 | 0.00001 | 0.00001 | 0.900209 | 0.00001 |
| 12_NB_M_EPK11 | 0.00001 | 0.00001 | 0.00001 | 0.00001 | 0.00001 | 0.99994 | 0.00001 |
| 13_NB_M_EPK17 | 0.00001 | 0.00001 | 0.00001 | 0.00001 | 0.00001 | 0.99994 | 0.00001 |
| 15_NB_F_EPK08 | 0.00001 | 0.00001 | 0.00001 | 0.00001 | 0.00001 | 0.99994 | 0.00001 |
| 16_NB_F_EPK02 | 0.226282 | 0.00001 | 0.00001 | 0.00001 | 0.00001 | 0.773668 | 0.00001 |
| 17_NB_M_EPK03 | 0.00001 | 0.00001 | 0.00001 | 0.00001 | 0.00001 | 0.99994 | 0.00001 |
| 23_NB_M_EPK01 | 0.00001 | 0.00001 | 0.00001 | 0.00001 | 0.00001 | 0.00001 | 0.99994 |
| 24_NB_M_EPK15 | 0.08412 | 0.00001 | 0.00001 | 0.00001 | 0.00001 | 0.00001 | 0.91583 |
| 25_NB_F_EPK10 | 0.00001 | 0.00001 | 0.00001 | 0.00001 | 0.00001 | 0.00001 | 0.99994 |
| 26_NB_F_EPK18 | 0.961073 | 0.00001 | 0.00001 | 0.038877 | 0.00001 | 0.00001 | 0.00001 |
| 30_NB_M_EPK09 | 0.828829 | 0.00001 | 0.00001 | 0.051866 | 0.00001 | 0.00001 | 0.119265 |
| 45_NB_F_EPK16 | 0.849131 | 0.00001 | 0.00001 | 0.028333 | 0.00001 | 0.00001 | 0.122495 |
| 49_NB_M_NUB13 | 0.471767 | 0.00001 | 0.00001 | 0.00001 | 0.00001 | 0.00001 | 0.528183 |
| JJK_4202 | 0.490042 | 0.00001 | 0.00001 | 0.00001 | 0.00001 | 0.00001 | 0.509908 |
| JJK_4205 | 0.489222 | 0.00001 | 0.00001 | 0.00001 | 0.00001 | 0.00001 | 0.510728 |
| JJK_4206 | 0.99994 | 0.00001 | 0.00001 | 0.00001 | 0.00001 | 0.00001 | 0.00001 |
| MSB_Mamm_228394 | 0.56624 | 0.00001 | 0.00001 | 0.00001 | 0.00001 | 0.43371 | 0.00001 |
| MSB_Mamm_228926 | 0.864387 | 0.00001 | 0.00001 | 0.00796 | 0.00001 | 0.00001 | 0.127613 |
| MSB_Mamm_228970 | 0.581215 | 0.00001 | 0.00001 | 0.418735 | 0.00001 | 0.00001 | 0.00001 |
| MSB_Mamm_229205 | 0.586581 | 0.00001 | 0.00001 | 0.413369 | 0.00001 | 0.00001 | 0.00001 |
| MSB_Mamm_229389 | 0.869032 | 0.00001 | 0.00001 | 0.002686 | 0.00001 | 0.00001 | 0.128242 |
| MSB_Mamm_229407 | 0.99994 | 0.00001 | 0.00001 | 0.00001 | 0.00001 | 0.00001 | 0.00001 |
| MSB_Mamm_229442 | 0.869992 | 0.00001 | 0.00001 | 0.001214 | 0.00001 | 0.00001 | 0.128753 |
| MSB_Mamm_229444 | 0.579085 | 0.00001 | 0.00001 | 0.420865 | 0.00001 | 0.00001 | 0.00001 |
| MSB_Mamm_229448 | 0.809083 | 0.00001 | 0.00001 | 0.047912 | 0.00001 | 0.00001 | 0.142965 |
| MSB_Mamm_229461 | 0.97032 | 0.00001 | 0.00001 | 0.02963 | 0.00001 | 0.00001 | 0.00001 |
| MSB_Mamm_229462 | 0.824518 | 0.00001 | 0.00001 | 0.0496 | 0.00001 | 0.00001 | 0.125841 |
| MSB_Mamm_229471 | 0.989448 | 0.00934 | 0.00001 | 0.00001 | 0.000011 | 0.00117 | 0.00001 |
| MSB_Mamm_229879 | 0.991872 | 0.008078 | 0.00001 | 0.00001 | 0.00001 | 0.00001 | 0.00001 |
| MSB_Mamm_72252 | 0.990333 | 0.008827 | 0.00001 | 0.00001 | 0.00001 | 0.0008 | 0.00001 |
| BSB32 | 0.995372 | 0.004578 | 0.00001 | 0.00001 | 0.00001 | 0.00001 | 0.00001 |
| FSB34 | 0.00001 | 0.00001 | 0.00001 | 0.00001 | 0.00001 | 0.00001 | 0.99994 |
| MSB_Mamm_156812 | 0.00001 | 0.00001 | 0.00001 | 0.00001 | 0.00001 | 0.00001 | 0.99994 |
| UAM_Mamm_71528 | 0.00001 | 0.00001 | 0.00001 | 0.00001 | 0.00001 | 0.00001 | 0.99994 |
| UAM_Mamm_71535 | 0.957529 | 0.00001 | 0.00001 | 0.042421 | 0.00001 | 0.00001 | 0.00001 |
| UAM_Mamm_71550 | 0.96805 | 0.00001 | 0.00001 | 0.0319 | 0.00001 | 0.00001 | 0.00001 |
| MSB_Mamm_149203 | 0.967239 | 0.00001 | 0.00001 | 0.032711 | 0.00001 | 0.00001 | 0.00001 |
| MSB_Mamm_149204 | 0.937087 | 0.00001 | 0.00001 | 0.062863 | 0.00001 | 0.00001 | 0.00001 |
| MSB_Mamm_149205 | 0.948875 | 0.00001 | 0.00001 | 0.051075 | 0.00001 | 0.00001 | 0.00001 |
| EPK001 | 0.95441 | 0.00001 | 0.00001 | 0.04554 | 0.00001 | 0.00001 | 0.00001 |
| EPK002 | 0.950592 | 0.00001 | 0.00001 | 0.049358 | 0.00001 | 0.00001 | 0.00001 |
| EPK003 | 0.964704 | 0.00001 | 0.00001 | 0.035246 | 0.00001 | 0.00001 | 0.00001 |
| EPK004 | 0.805111 | 0.00001 | 0.00001 | 0.064946 | 0.00001 | 0.00001 | 0.129903 |
| EPK005 | 0.953487 | 0.00001 | 0.00001 | 0.046463 | 0.00001 | 0.00001 | 0.00001 |
| EPK006 | 0.00001 | 0.00001 | 0.99994 | 0.00001 | 0.00001 | 0.00001 | 0.00001 |
| EPK007 | 0.00001 | 0.00001 | 0.99994 | 0.00001 | 0.00001 | 0.00001 | 0.00001 |
| EPK008 | 0.00001 | 0.00001 | 0.00001 | 0.99994 | 0.00001 | 0.00001 | 0.00001 |
| EPK010 | 0.00001 | 0.00001 | 0.00001 | 0.99994 | 0.00001 | 0.00001 | 0.00001 |
| EPK011 | 0.00001 | 0.00001 | 0.00001 | 0.99994 | 0.00001 | 0.00001 | 0.00001 |
| EPK012 | 0.00001 | 0.00001 | 0.00001 | 0.99994 | 0.00001 | 0.00001 | 0.00001 |
| EPK014 | 0.00001 | 0.00001 | 0.00001 | 0.99994 | 0.00001 | 0.00001 | 0.00001 |
| EPK017 | 0.00001 | 0.00001 | 0.00001 | 0.99994 | 0.00001 | 0.00001 | 0.00001 |
| EPK018 | 0.00001 | 0.00001 | 0.00001 | 0.99994 | 0.00001 | 0.00001 | 0.00001 |
| EPK019 | 0.00001 | 0.00001 | 0.00001 | 0.99994 | 0.00001 | 0.00001 | 0.00001 |
| MSB_Mamm_228743 | 0.00001 | 0.00001 | 0.00001 | 0.99994 | 0.00001 | 0.00001 | 0.00001 |
| MSB_Mamm_228765 | 0.00001 | 0.00001 | 0.00001 | 0.99994 | 0.00001 | 0.00001 | 0.00001 |
| MSB_Mamm_228782 | 0.00001 | 0.00001 | 0.00001 | 0.99994 | 0.00001 | 0.00001 | 0.00001 |
| MT_10_56 | 0.00001 | 0.00001 | 0.00001 | 0.99994 | 0.00001 | 0.00001 | 0.00001 |
| MT_14_23 | 0.00001 | 0.00001 | 0.00001 | 0.99994 | 0.00001 | 0.00001 | 0.00001 |
| JKK_4190 | 0.00001 | 0.00001 | 0.00001 | 0.99994 | 0.00001 | 0.00001 | 0.00001 |
| JKK_4191 | 0.00001 | 0.00001 | 0.00001 | 0.99994 | 0.00001 | 0.00001 | 0.00001 |
| JKK_4192 | 0.00001 | 0.00001 | 0.00001 | 0.99994 | 0.00001 | 0.00001 | 0.00001 |
| MZ10760 | 0.00001 | 0.00001 | 0.00001 | 0.99994 | 0.00001 | 0.00001 | 0.00001 |
| MZ10762 | 0.99994 | 0.00001 | 0.00001 | 0.00001 | 0.00001 | 0.00001 | 0.00001 |
| MZ10764 | 0.00001 | 0.00002 | 0.00001 | 0.00001 | 0.99993 | 0.00001 | 0.00001 |
| MZ10770 | 0.00001 | 0.00001 | 0.00001 | 0.020885 | 0.979065 | 0.00001 | 0.00001 |
| MZ10772 | 0.00001 | 0.00001 | 0.000019 | 0.000013 | 0.999928 | 0.00001 | 0.00001 |
| MZ10774 | 0.99994 | 0.00001 | 0.00001 | 0.00001 | 0.00001 | 0.00001 | 0.00001 |
| MZ10804 | 0.99994 | 0.00001 | 0.00001 | 0.00001 | 0.00001 | 0.00001 | 0.00001 |
| MZ10808 | 0.99994 | 0.00001 | 0.00001 | 0.00001 | 0.00001 | 0.00001 | 0.00001 |
| MZ10812 | 0.99994 | 0.00001 | 0.00001 | 0.00001 | 0.00001 | 0.00001 | 0.00001 |
| MZ10813 | 0.99994 | 0.00001 | 0.00001 | 0.00001 | 0.00001 | 0.00001 | 0.00001 |
| MZ10814 | 0.999934 | 0.00001 | 0.00001 | 0.00001 | 0.000016 | 0.00001 | 0.00001 |
| MZ10815 | 0.99994 | 0.00001 | 0.00001 | 0.00001 | 0.00001 | 0.00001 | 0.00001 |
| MZ10884 | 0.99994 | 0.00001 | 0.00001 | 0.00001 | 0.00001 | 0.00001 | 0.00001 |
| MZ10885 | 0.99994 | 0.00001 | 0.00001 | 0.00001 | 0.00001 | 0.00001 | 0.00001 |
| MZ11010 | 0.99994 | 0.00001 | 0.00001 | 0.00001 | 0.00001 | 0.00001 | 0.00001 |
| MZ11011 | 0.99994 | 0.00001 | 0.00001 | 0.00001 | 0.00001 | 0.00001 | 0.00001 |
| MZ11012 | 0.99994 | 0.00001 | 0.00001 | 0.00001 | 0.00001 | 0.00001 | 0.00001 |
| MSB_Mamm_48208 | 0.00001 | 0.00001 | 0.00001 | 0.00001 | 0.99994 | 0.00001 | 0.00001 |
| MSB_Mamm_48209 | 0.00001 | 0.00001 | 0.00001 | 0.00001 | 0.99994 | 0.00001 | 0.00001 |
| MSB_Mamm_48210 | 0.00001 | 0.00001 | 0.00001 | 0.00001 | 0.99994 | 0.00001 | 0.00001 |

**Table S5.** D-statistic for the *Peromyscus maniculatus* species complex.

| P1 | P2 | P3 | Dstatistic | Z.score | p.value | f4.ratio | BBAA | ABBA | BABA | BH.pval |
| --- | --- | --- | --- | --- | --- | --- | --- | --- | --- | --- |
| maniculatus | midwest | keeni | 0.0505232 | 3.6215 | 0.00029291 | 0.0705548 | 1190.06 | 1039.05 | 939.111 | 0.00073226 |
| keeni | midwest | melanotis | 0.0119182 | 0.77344 | 0.439262 | 0.00267133 | 2353.87 | 644.859 | 629.669 | 0.62751714 |
| polionotus | midwest | keeni | 0.0858516 | 5.61302 | 1.99E-08 | 0.117443 | 1134.69 | 1102.09 | 927.816 | 6.63E-08 |
| midwest | sonoriensis | keeni | 0.0145406 | 1.63196 | 0.102687 | 0.022 | 1150.97 | 1014.56 | 985.477 | 0.205374 |
| maniculatus | midwest | melanotis | 0.00634668 | 0.390536 | 0.69614 | 0.0013262 | 2457.85 | 597.109 | 589.578 | 0.790291 |
| polionotus | maniculatus | midwest | 0.0811105 | 8.43135 | 2.30E-16 | 0.0821701 | 1608.03 | 788.792 | 670.433 | 1.15E-15 |
| sonoriensis | midwest | maniculatus | 0.0845351 | 13.3754 | 2.30E-16 | 0.121569 | 1128.37 | 1113.88 | 940.237 | 1.15E-15 |
| polionotus | midwest | melanotis | 0.0196768 | 0.877235 | 0.380359 | 0.00422734 | 2368 | 623.964 | 599.882 | 0.58516769 |
| sonoriensis | midwest | melanotis | 0.0033049 | 0.319385 | 0.749434 | 0.0007148 | 2489.48 | 615.786 | 611.73 | 0.790291 |
| sonoriensis | midwest | polionotus | 0.0916689 | 11.7135 | 2.30E-16 | 0.0472944 | 1211.34 | 1078.39 | 897.286 | 1.15E-15 |
| keeni | maniculatus | melanotis | 0.00585102 | 0.322882 | 0.746785 | 0.00134684 | 2277.25 | 658.298 | 650.639 | 0.790291 |
| polionotus | maniculatus | keeni | 0.0530946 | 4.68246 | 2.83E-06 | 0.0497555 | 1802.82 | 732.189 | 658.358 | 8.10E-06 |
| keeni | sonoriensis | maniculatus | 0.0374819 | 5.76695 | 8.07E-09 | 0.0513455 | 1121.59 | 1069.87 | 992.567 | 3.23E-08 |
| polionotus | keeni | melanotis | 0.00659808 | 0.316363 | 0.751727 | 0.001545 | 2225.46 | 671.353 | 662.552 | 0.790291 |
| keeni | sonoriensis | melanotis | 0.00866466 | 0.630784 | 0.528182 | 0.0019579 | 2388.63 | 648.022 | 636.889 | 0.70424267 |
| keeni | sonoriensis | polionotus | 0.0128861 | 1.98588 | 0.0470471 | 0.00668649 | 1189.92 | 1012.55 | 986.791 | 0.10454911 |
| polionotus | maniculatus | melanotis | 0.018122 | 0.961641 | 0.33623 | 0.0028901 | 3144.9 | 462.478 | 446.014 | 0.57602167 |
| maniculatus | sonoriensis | melanotis | 0.00273972 | 0.265933 | 0.790291 | 0.00061185 | 2327.52 | 635.872 | 632.397 | 0.790291 |
| polionotus | maniculatus | sonoriensis | 0.0893864 | 9.22413 | 2.30E-16 | 0.317128 | 1758.25 | 765.1 | 639.544 | 1.15E-15 |
| polionotus | sonoriensis | melanotis | 0.0151463 | 0.943132 | 0.345613 | 0.00350101 | 2235.98 | 668.344 | 648.4 | 0.57602167 |

**Table S6.** Top five ENMs for Peromyscus maniculatus throughout its known range in North America.

| fc | rm | auc.val.avg | or.mtp.avg | or.mtp.sd |
| --- | --- | --- | --- | --- |
| LQ | 2 | 0.71414052 | 0.001 | 0.002 |
| LQ | 3 | 0.71348088 | 0.001 | 0.002 |
| LQ | 4 | 0.71233866 | 0.001 | 0.002 |
| LQ | 1 | 0.71088924 | 0.001 | 0.002 |
| LQ | 5 | 0.70700971 | 0.001 | 0.002 |

**Table S7.** Summary of traits included in final phenotypic analyses. First column denotes full name, and second column the shorthand labels used in text and supplementary figures. Third column defines trait as measured in this study.

| Trait | Shorthand | Definition |
| --- | --- | --- |
| Caudal vertebrae 1-31 | c1-c31 | Lengths of the caudal vertebrae |
| Final caudal vertebrae length | tv_final | Length of last measurable caudal vertebrae |
| Caudal vertebrae number | max_tail_vert | No. caudal vertebrae varies by individual |
| Longest caudal vertebrae | tv_longest | Length of largest caudal vertebra (c1-c31) |
| Sacral vertebrae 1-4 | s1-s4 | All individuals have 4 sacral vertebrae |
| Weight | weight | Weight in grams |
| Total length | total | Length of individual from nose to tail tip |
| Body length | body | Total length minus the tail |
| Tail/body ratio | tail_body | Length from base to tip, scaled by body size |
| Ear/body ratio | ear_body | Length from notch (base) to tip, scaled by body size |
| Foot/body ratio | foot_body | Corresponds to hind foot, measured from heel to toe, scaled by<br>body_size |
| Whisker lengths | (a,b,g,d)_body | a, b, g, and d denote alpha, beta, gamma, and delta caudal offset<br>whiskers, scaled by body size |
| Skull length | skull | Length from base of skull to nose, measured by X-ray |
| Pelvis length | pelvis_length | Length of pelvis |
| Ilium width | ilium_width | Length of ilium |
| Ischium width | ischium_width | Length of ischium |

**Table S8.**
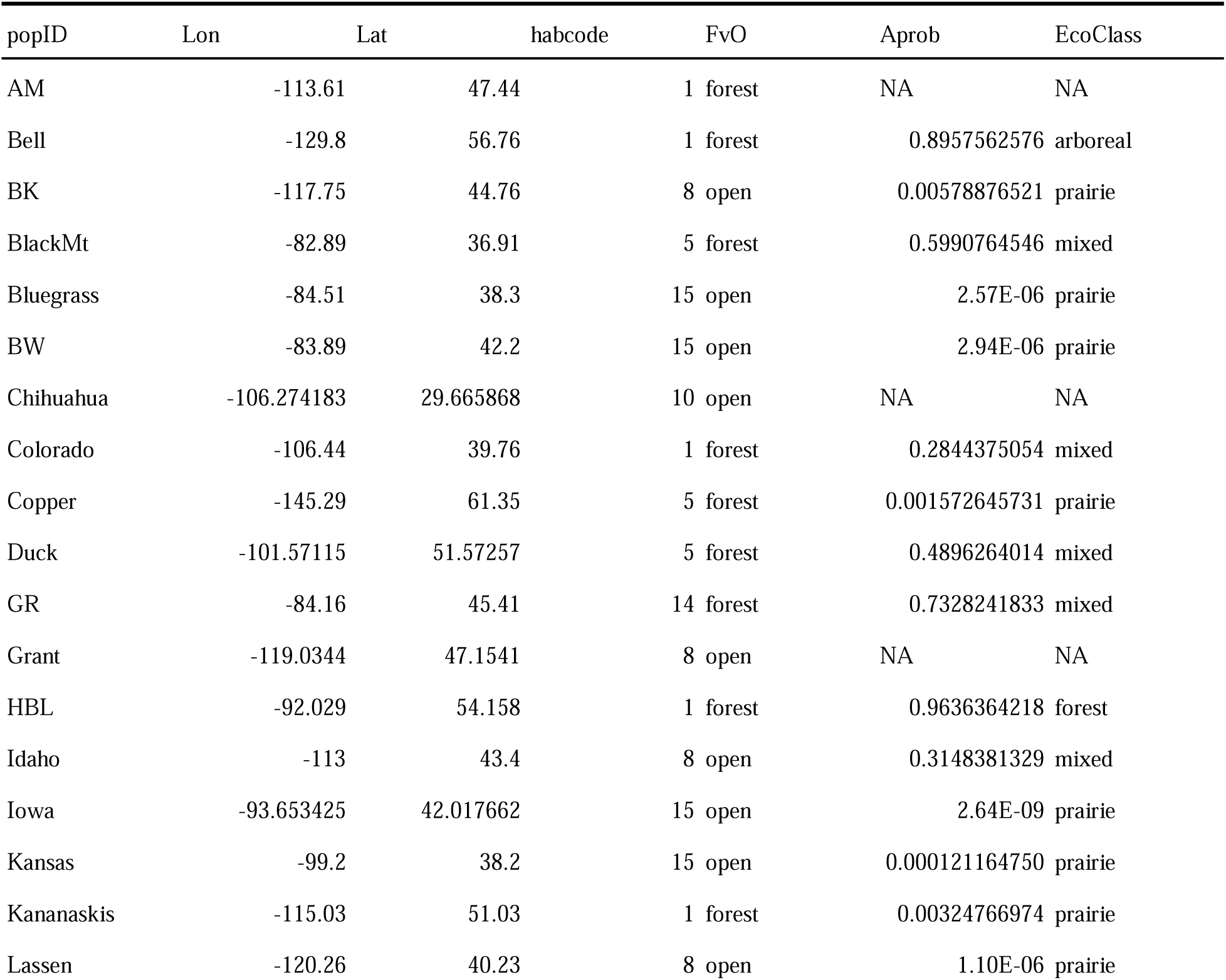
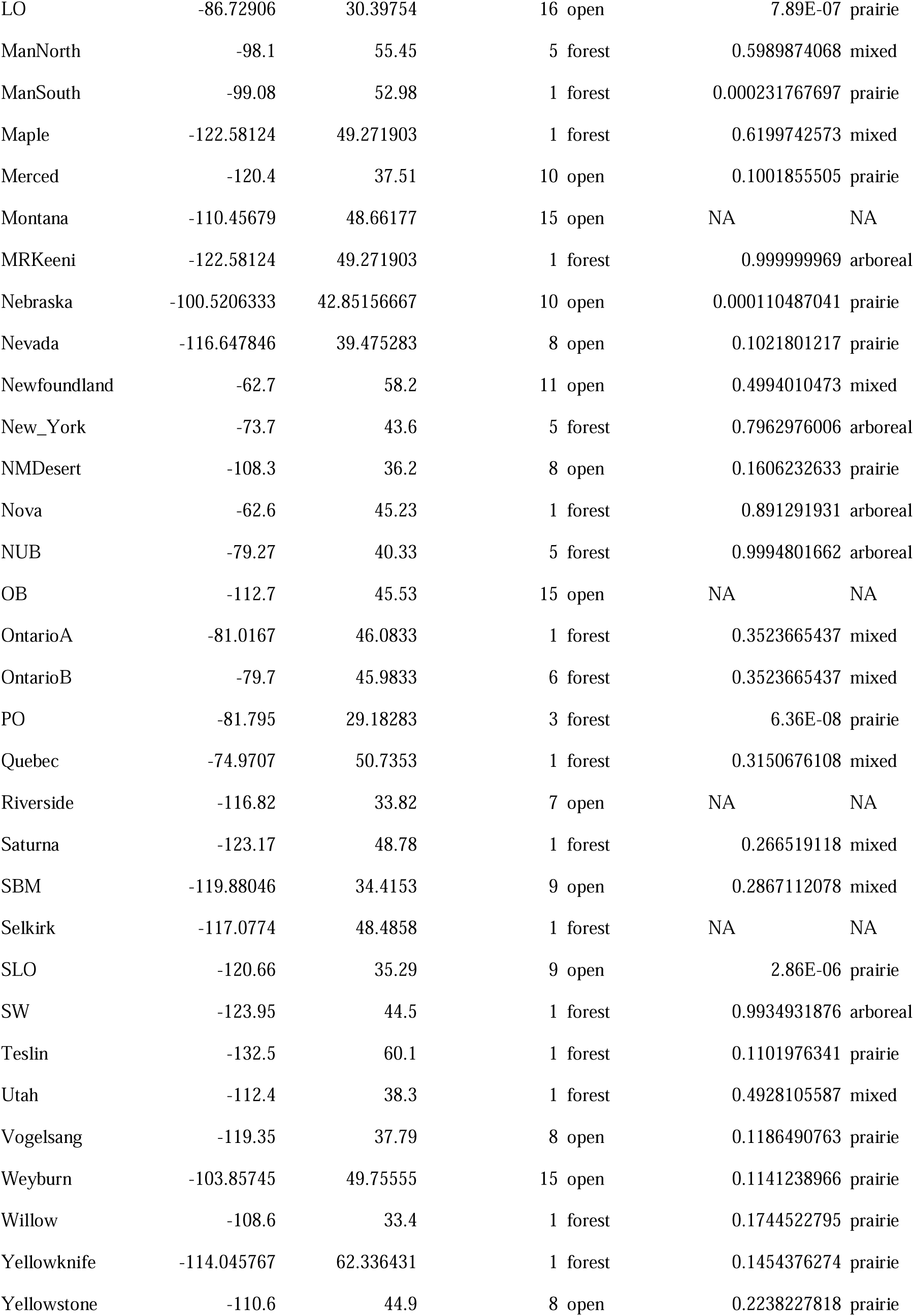
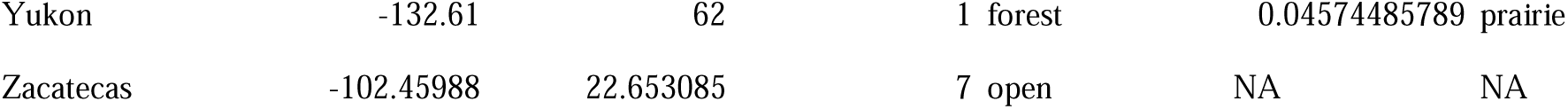
Habitat and ecotype classifications. All populations with genomic data are included in this table, which directly corresponds to the results presented in Fig. 4. <u>popID</u>, <u>Lon</u>, and <u>Lat</u>, are the same as in Extended Data Table 2. <u>Habcode</u> = Most common habitat type following NALCMS classification, and <u>FvO</u> = Forest v. Open habitat classification to <u>Habcode</u> (see Methods section). <u>Aprob</u> = Posterior probability of the population being classified as the arboreal (1) or prairie (0) ecotype. <u>EcoClass</u> = Qualitative classification of habitat type based on <u>Aprob</u>, where <u>Aprob</u> <0.25 = Prairie, <u>Aprob</u> in [0.25,0.75] = Mixed, and <u>Aprob</u> > 0.75 = Arboreal.

**Figure S1.**
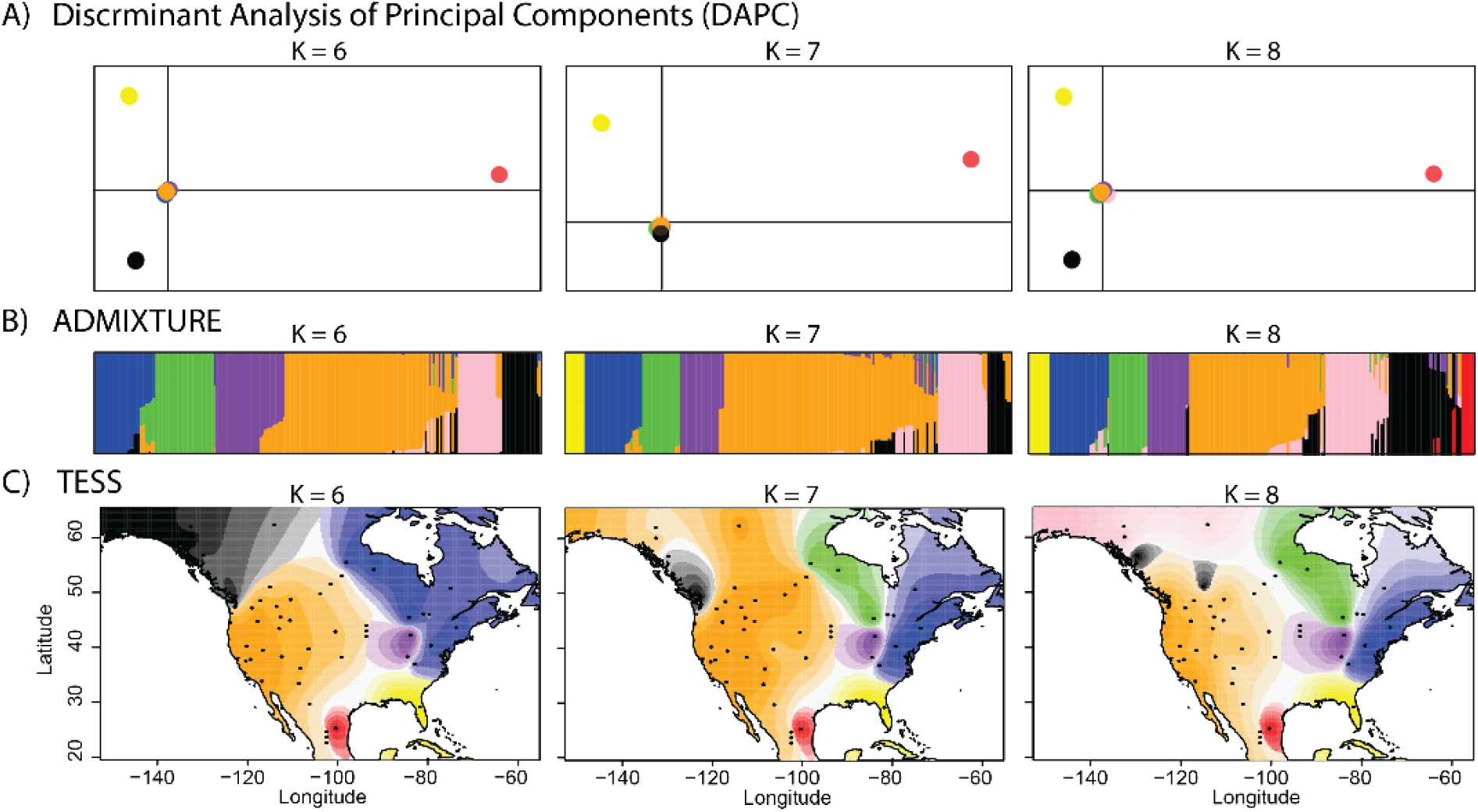
**A)** Discriminant analysis of principal components (DAPC), **B)** ADMIXTURE analysis, and **C)** TESS3R analysis of genome wide SNPS for K=6 through K=8.

**Figure S2.**
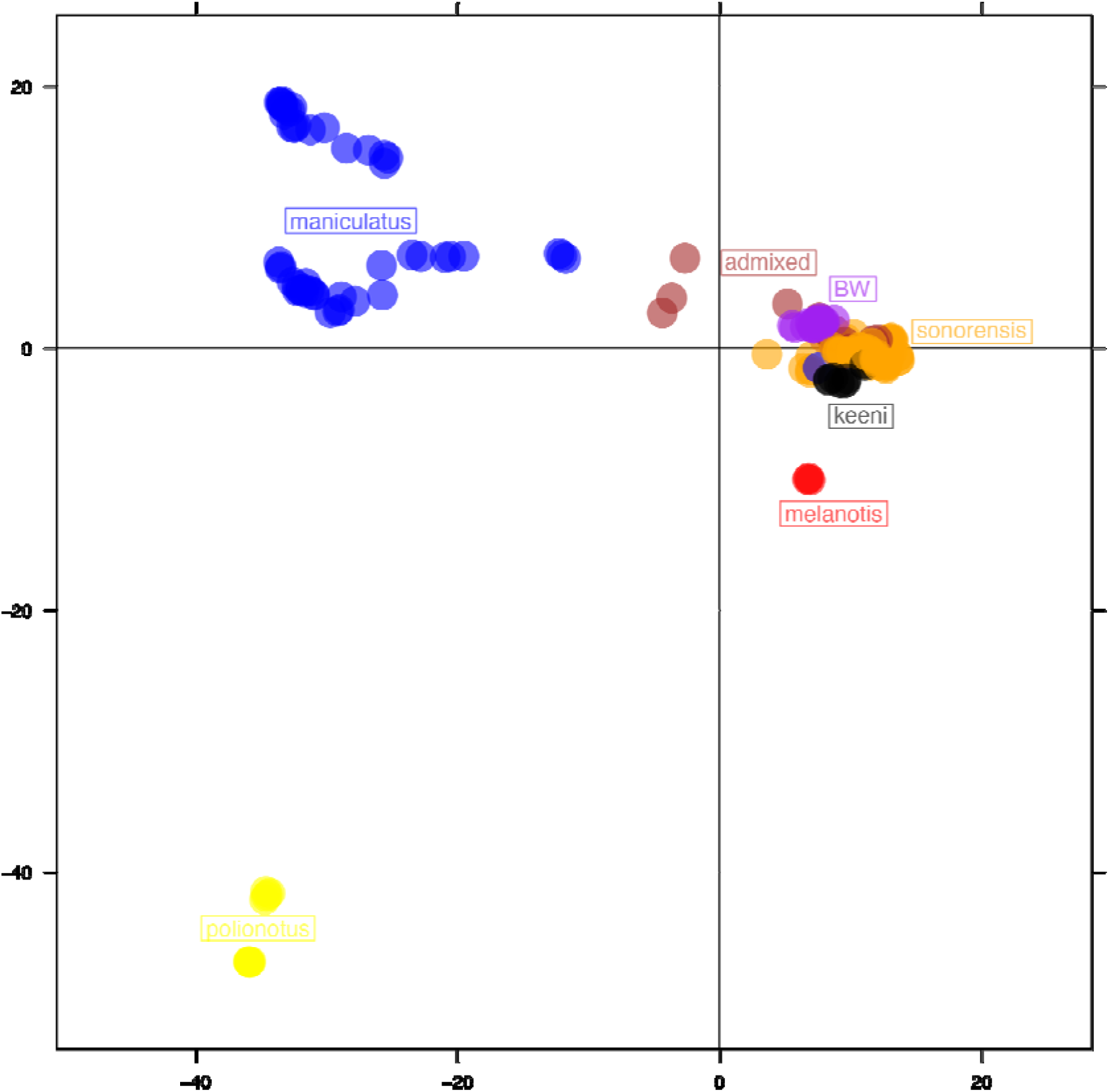
Principal components analysis indicating admixed individuals for the *Peromyscus* species complex.

**Figure S3.**
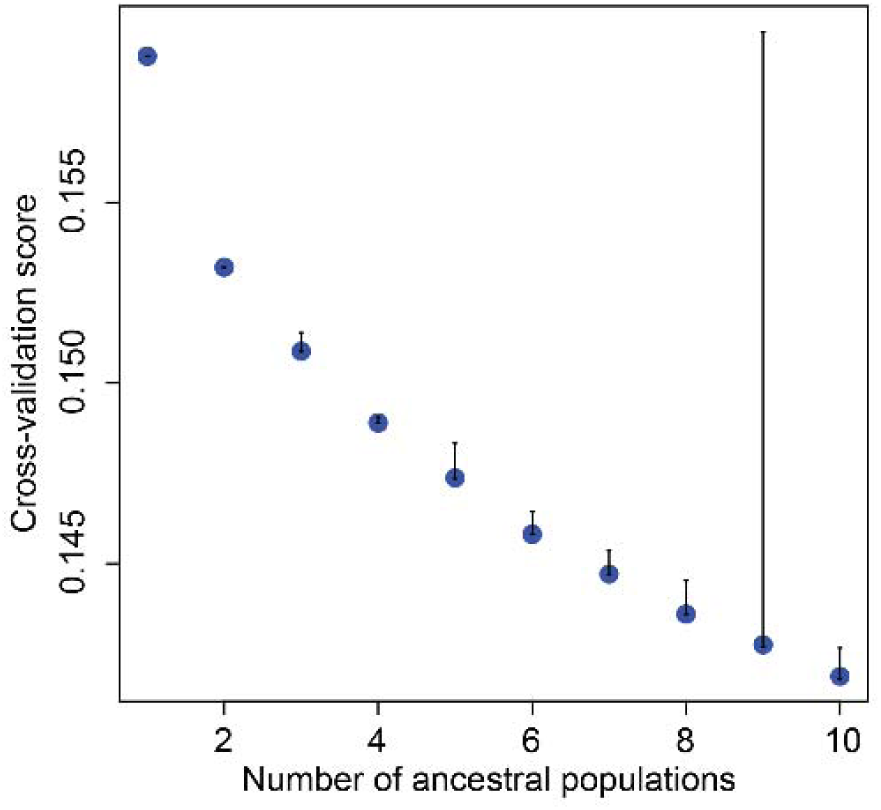
The cross validation results from *tess3r* on *Peromyscus maniculatus* across its range in North America.

**Figure S4.**
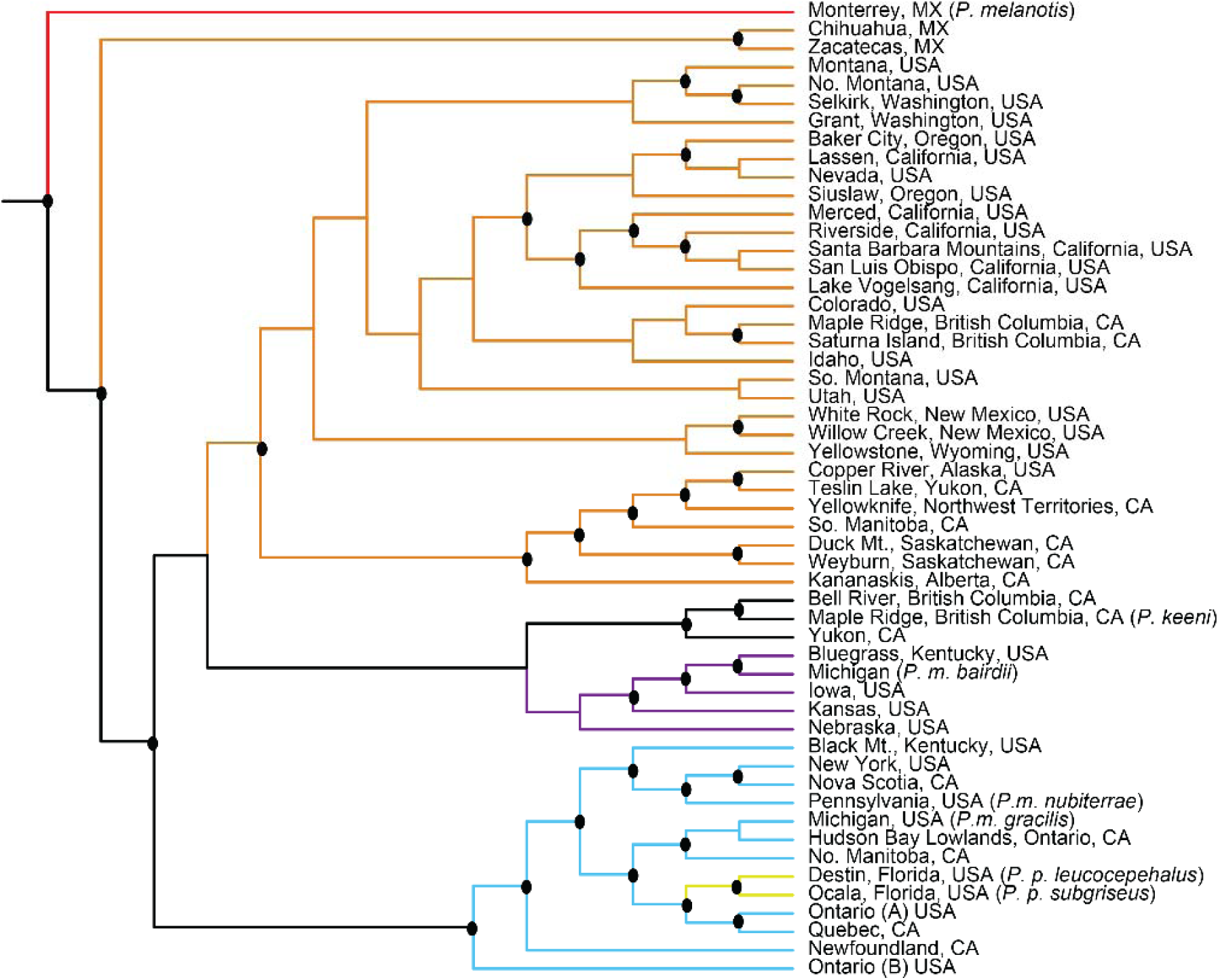
The phylogenetic relationships of populations within the *Peromyscus maniculatus* species complex across its range in North America using SVDquartets for the “whole” dataset. The black circles at nodes indicate bootstrap support greater than 70.

**Figure S5.**
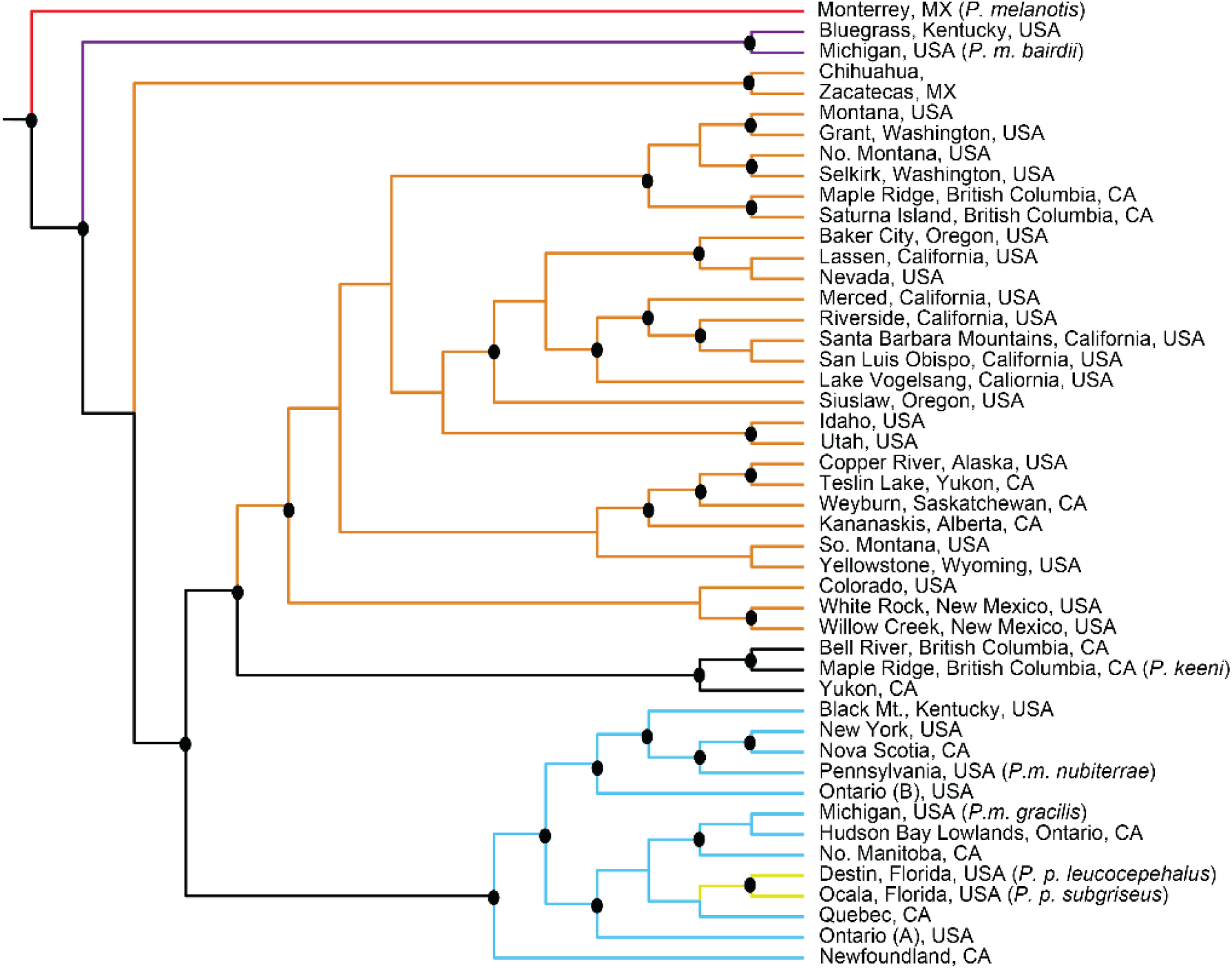
The phylogenetic relationships of populations within the *Peromyscus maniculatus* species complex across its range in North America using SVDquartets for the “reduced” dataset. The black circles at nodes indicate bootstrap support greater than 70.

**Figure S6.**
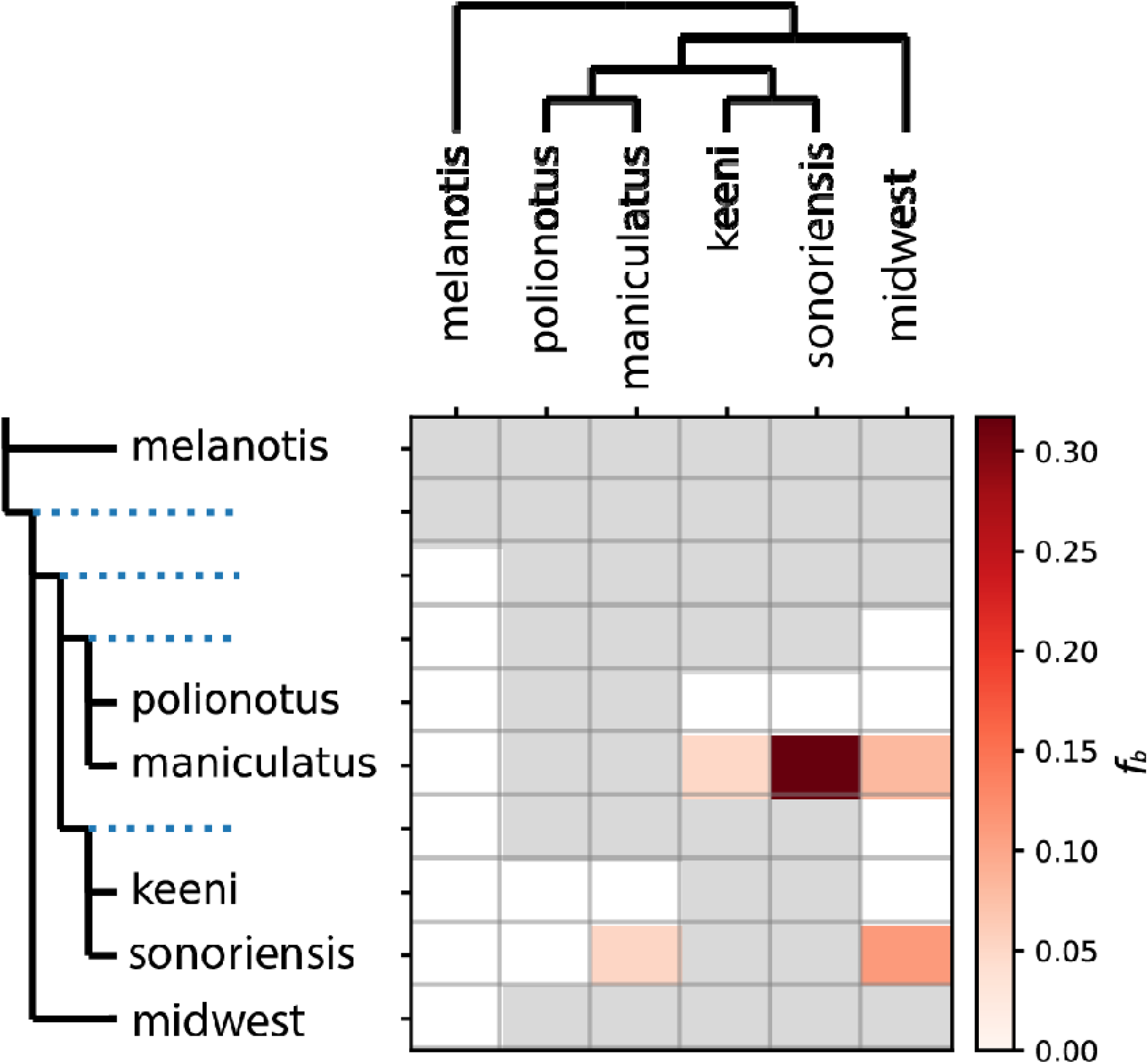
F-branch statistics for the *Peromyscus maniculatus* species complex across its range in North America using *Dsuite*.

**Figure S7.**
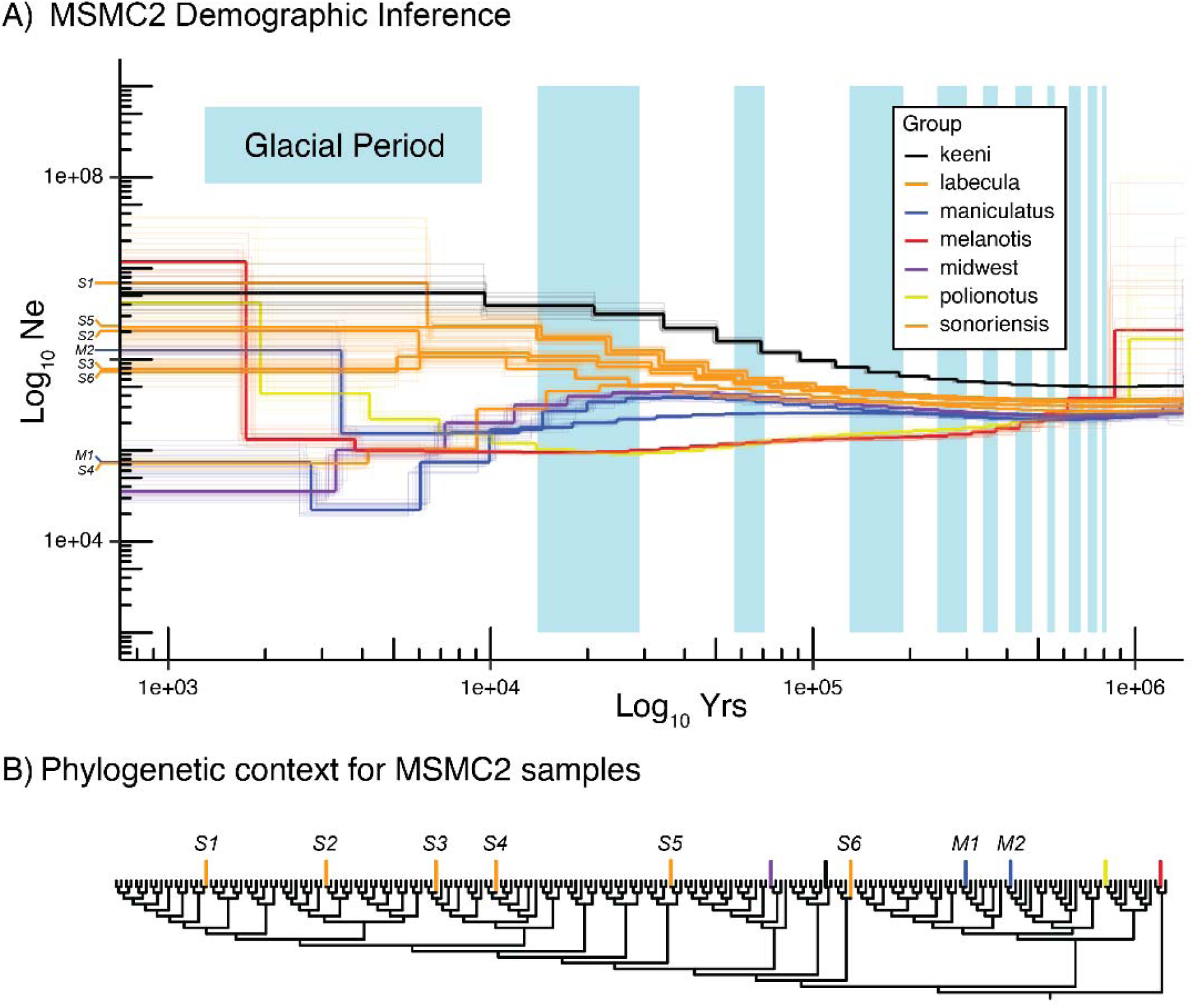
MSMC2 population size histories. **A)** N_e_ through time for individuals selected from the main clades of the *P. maniculatus* species complex. Color coding corresponds to phylogenetic color coding in Figure 2. X-axis has been transformed to years assuming a *de novo* mutation rate of 5.34e-9 and generation time of 0.5 years. **B)** Phylogeny from Fig. 2 recreated here to highlight the samples selected for demographic reconstruction. *Sonoriensis* and *maniculatus* have additional labels (i.e. S1, M1) t distinguish different individuals in A) and B).

**Figure S8.**
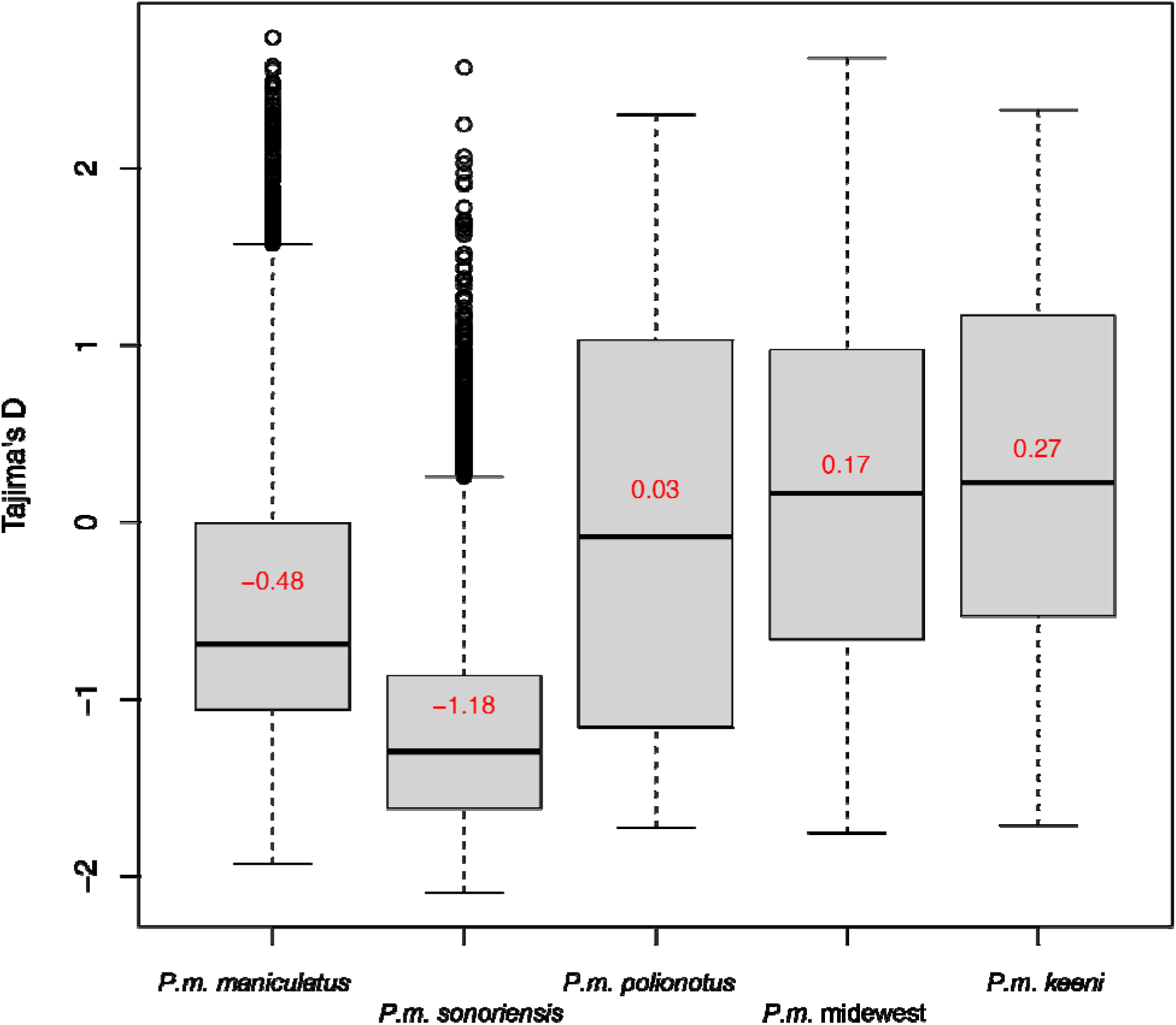
Tajima’s D for the five lineages within the *Peromyscus maniculatus* species complex across its range in North America.

**Figure S9.**
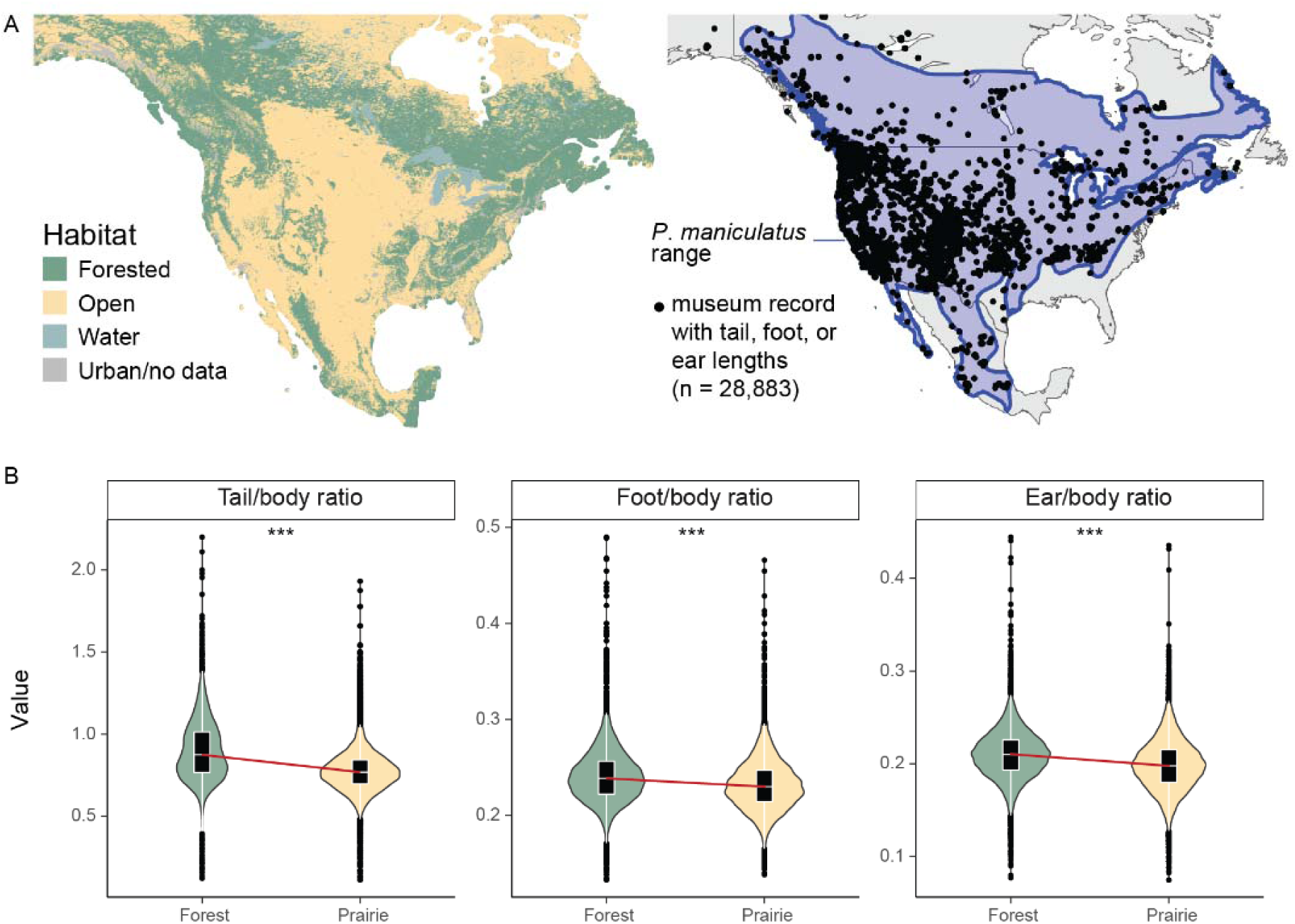
Associations between three traits and habitat type based on museum records. **A)** Left: map of North America color coded by “forest” or “prairie” habitat as summarized at 3km resolution from NALCMS satellite data. Right: estimated range of *P. maniculatus*, and distribution of museum record containing any phenotypic information on tail, foot, or ear lengths. **B)** Phenotype-habitat associations as determined from *P. maniculatus* museum records. Tail, foot, and ear lengths are all scaled by body size. ‘***’ denotes p<2.2e-16, as determined by two-sided t-tests.

**Figure S10.**
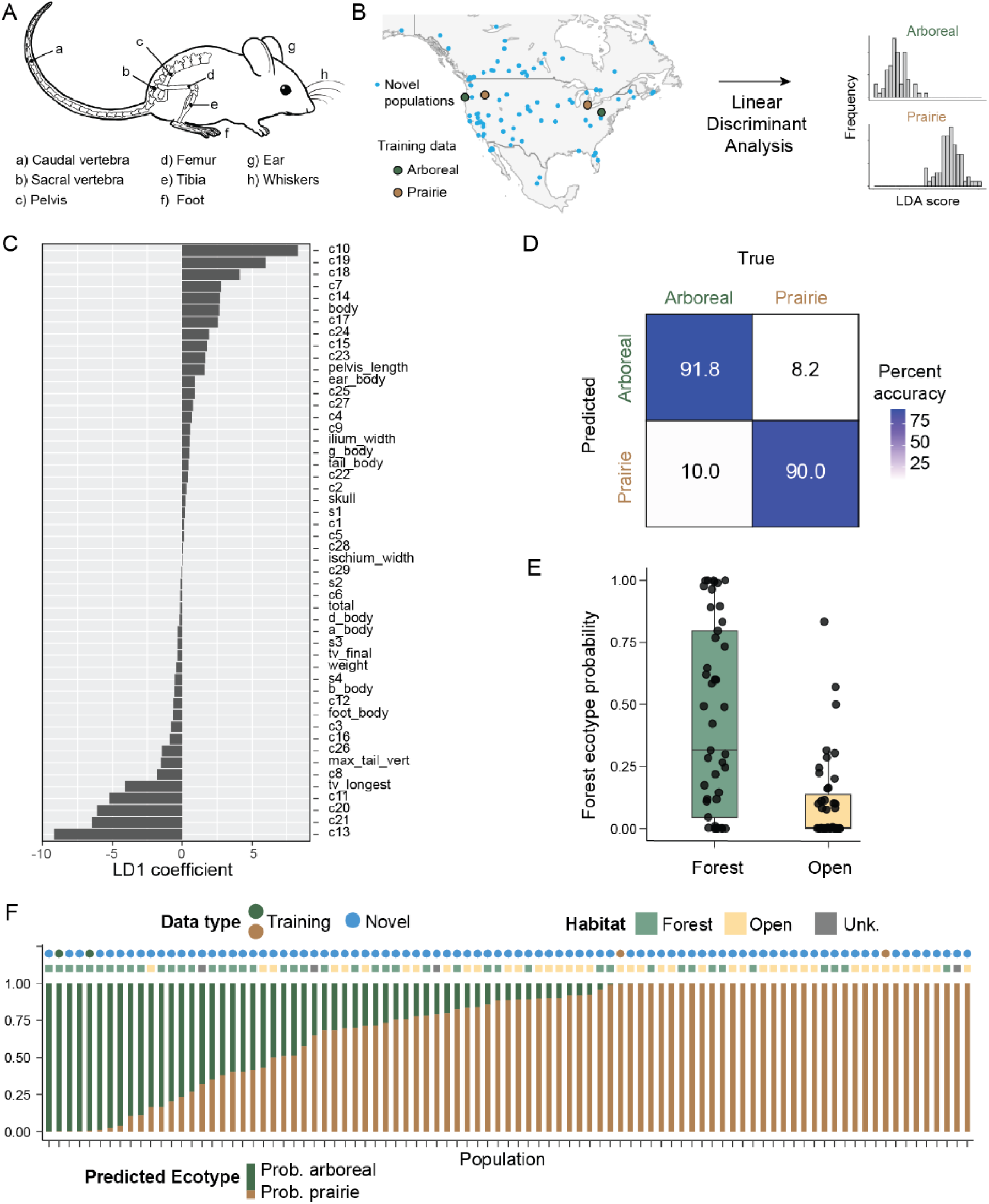
Identifying arboreal and prairie ecotypes across North America. **A)** Simplified diagram of a deer mouse displaying morphological areas of interest. For brevity, only general anatomical areas are annotated, not the full set of measured traits (see Table S1 for full set). **B)** Map of North America with locations of known arboreal and prairie ecotypes, as well as previously uncharacterized populations. Histograms show the ability of linear discriminant analysis (LDA) to discriminate between arboreal and prairie individuals in training data. **C)** LD1 trait coefficients for model based on training forest-prairie data. Variables are labeled using shorthand (Table S1) **D)** Confusion matrix showing the results of leave-one-out-cross-validation on training forest-prairie data. **E)** Association between ecotype and habitat type. Boxes summarize FEPP distributions for populations by habitat, displaying median, first quartile (Q1), first quartile (Q3) and 1.5*inter-quartile range (IQR) **F)** Ecotype predictions of novel populations by linear discriminant model, and correspondence to habitat type. Y-axis denotes mean posterior probability of ecotype across all samples in a population.

## Extended Data

The following are captions for extended data files that did not fit within the main text or supplementary materials of this paper. These files are archived on Zenodo (10.5281/zenodo.18227645).

**Extended Data Table 1.** List of individual samples used in this study. <u>Species</u> = Purported species as original identified by collector or institution. <u>Institution/Source</u> = Origin of specimen. <u>SampleID</u> = Sample name as referenced in text and on NCBI SRA where applicable.. <u>popID</u> = Population shorthand name (see Extended Data Table 2). <u>accessionID</u> = Museum accession number, if applicable. <u>Sequenced</u> = Indicates whether the individual was sequenced (Y) or not (N).

**Extended Data Table 2.** All population names, shorthand names, and coordinates used in this study, either for phenotyping and/or genotyping. <u>Population</u> = Population name. <u>popID</u> = Shorthand name used throughout the manuscript and analyses. <u>Lat</u> = latitude. <u>Lon</u> = longitude.

**Extended Data Table 3.** Geographic coordinates used in ENMs for Peromyscus maniculatus throughout its known range in North America. <u>Lat</u> = latitude. <u>Lon</u> = longitude.

**Extended Data Table 4.** Spatially thinned geographic coordinates used in ENMs for Peromyscus maniculatus throughout its known range in North America. <u>Lat</u> = latitude. <u>Lon</u> = longitude.

**Extended Data Video 1.** Ecological niche models of P. maniculatus throughout its range in North America over the last 21 thousand years, at 500 year intervals.

**Extended Data Files** - Phylogenetic reconstructions at the individual level:

1) **sample_ALL.tree** (all samples)
2) **sample_NOADMIX.tree** (removed admixed individuals >.70 admixture scores
3) **pop_ALL.tree** (all samples)
4) **pop_NOADMIX.tree** (removed admixed individuals >.70 admixture scores)

## Data Availability

All original code will be made available on Github (https://github.com/twooldridge/PeromyscusPhyloGeo) upon formal publication. All sequence data will be deposited on the NCBI SRA database.

